# Host tRNA modifications drive efficient translation of influenza A virus genome and impact host antiviral stress responses

**DOI:** 10.64898/2026.04.21.719891

**Authors:** Diana Roberta Ribeiro, Alexandre Nunes, Mie Monti, Laia Llovera, Maximilian Berg, Kira Kerkhoff, Marisa Pereira, Ana Rita Guimarães, Stefanie Kaiser, Eva Maria Novoa, Daniela Ribeiro, Ana Raquel Soares

## Abstract

Influenza A virus (IAV), as all other viruses, is completely dependent on the host translation machinery components, including host transfer RNAs (tRNAs), to effectively decode its genome. However, while the human genome is biased towards cytosine (C) and guanosine (G)-ending codons, the IAV genome is skewed towards adenine (A) and uridine (U)-ending codons. Nevertheless, translation of IAV’s RNA genome is highly efficient. Here we show that host tRNA and tRNA epitranscriptome dynamics are important regulators of IAV RNA translation and host antiviral responses. We show that the levels of several tRNA modifications, including 5-methylcarboxymethyluridine (mcm^5^U_34_) and 5-methoxycarbonylmethyl-2-thiouridine (mcm^5^s^2^U_34_), and their cognate writers, vary over the course of IAV infection. Additionally, we demonstrate that a set of tRNAs are preferentially recruited to ribosomes upon IAV infection, in line with IAV codon usage requirements. We further show that loss of ELP3, the catalytic subunit of the elongator complex, which is involved in the catalysis of mcm^5^U_34_ and of mcm^5^s^2^U_34_, induces tRNA hypomodifications, impairs translation of codon biased IAV genes and triggers the integrated stress response (ISR), interfering with IAV propagation. Taken together, our results uncover the relevance of host tRNAs and their modifications for optimal expression of viral genomes and host antiviral responses, setting the tRNA epitranscriptome as a promising target for the development of host-based antiviral therapies.

## Introduction

Although genetically diverse, all viruses rely on the host cellular machinery to efficiently replicate. Among other cellular components, viruses hijack the host translation machinery, including the host transfer RNAs (tRNAs), to synthesize their own proteins at high rates^1^. However, the mechanisms by which this is done are still not completely clear, as host tRNA pools are not naturally adapted to efficiently translate most viral RNA genomes^1^. As the effector molecules of translation, tRNAs recognize messenger RNA (mRNA) codons through their anticodons, allowing decoding and incorporation of the corresponding amino acids into the nascent polypeptides^2^. The rate at which a given mRNA codon is decoded depends on factors such as tRNA availability, codon-anticodon interactions and tRNA chemical modifications^2–4^. Cellular tRNA concentrations and tRNA pools typically reflect the cellular codon usage, ensuring a balance between tRNA supply and demand. This balance contributes to translation efficiency and fidelity, ensuring cellular proteome homeostasis according to the cellular needs^5,6^. tRNA modifications are another important factor for efficient codon recognition and reading frame maintenance, especially when occurring at or near the tRNA anticodon region^7–10^. These modifications are highly dynamic across cells and tissues and can rapidly change to reprogram translation under specific stress conditions^6,11,12^, constituting a post-transcriptional regulatory layer. Collectively, tRNA modifications are known as the tRNA epitranscriptome and are catalyzed by different classes of tRNA modifying enzymes^2^. These are often disrupted or mutated in several diseases, ranging from neurological disorders^13–15^ to cancer^16–19^. However, the relevance of the tRNA epitranscriptome in the context of infection is only now beginning to emerge^20,21^.

In humans and other species that are natural influenza A virus (IAV) hosts, mRNAs are enriched in guanosine- (G) or cytosine- (C) ending codons. On the other hand, IAV’s RNA genome is biased towards adenosine- (A) or uridine- (U) ending codons, for which cognate host tRNAs are scarce and are generally highly modified at their anticodons to efficiently recognize the corresponding mRNA codons^22–24^. Nonetheless, translation of IAV transcripts is highly efficient, suggesting that IAV has evolved specific mechanisms to adjust the host tRNA supply in accordance with its own mRNA repertoire^22,24^. One of the few studies on this matter found that, during IAV RNA translation in host cells, tRNAs matching the viral RNA codon bias are preferentially associated with polysomes, despite the total tRNA pool remaining unchanged, suggesting the existence of local tRNA pools at sites of viral translation^25^. Nevertheless, the effects of IAV infection on the levels and dynamics of tRNA modifications remain unexplored.

In recent years, evidence of host tRNA modification dynamics in response to RNA viruses’ infection has been reported. For example, the dengue (DENV), chikungunya (CHIKV), severe acute respiratory syndrome coronavirus 2 (SARS-Cov-2) and Zika (ZIKV) viruses were recently shown to reprogram codon optimality through the host tRNA epitranscriptome^26–29^. Early DENV infection was shown to reduce ALKBH1 and its dependent f^5^Cm modification in tRNA^Leu^_CAA_ to enhance the decoding of transcripts rich in the non-cognate UUG codon^26^. Remarkably, the DENV NS5 protein acted as a tRNA-modifying enzyme at later stages of infection, introducing f^5^Cm into the above-mentioned tRNA to preferentially translate proviral UUA-rich mRNAs^26^. CHIKV was shown to increase 5-methylcarboxymethyluridine (mcm^5^U_34_) wobble base modifications by increasing the expression of KIAA1456 and hence facilitate the codon-biased decoding of the CHIKV A/U-ending codons^27^. Also SARS-CoV2 and ZIKV infections were shown to induce ncm^5^U, mcm^5^U and mcm^5^s^2^U hypermodification at wobble uridines of tRNAs, after several infection cycles in different cell lines^27–29^. Recent data have also shed light on the importance of tRNA modifications in the context of the cellular immune response. For instance, the levels of the tRNA-modifying enzymes TRMT61A and TRMT6 were increased to catalyze specific tRNA methylations during the activation of naïve T cells^30^. On the other hand, at the peak of T cell proliferation, the levels of yW and ms^2^t^6^A were drastically decreased to possibly promote ribosomal frameshifting, which may explain the preference of HIV-1 for these cell types^31^. Also, 2’-O-methylthymidine modification at position 54 of tRNAs were reported to inhibit immune stimulation and reduce the immune response in PBMCs^32^. Collectively, these results indicate that the reprogramming of tRNA modifications may also be correlated with coordination of host antiviral responses. Here we show that a single cycle of IAV infection induces dynamic changes in host U34 tRNA modifications and cognate tRNA modifying enzyme levels, without affecting total tRNA pools. We also show that specific tRNAs, namely tRNA^Lys^_UUU_ and tRNA^Glu^_UUC_ that bear U34 modifications, are enriched in translating ribosomes at time points of the infection cycle coinciding with viral RNA translation. Additionally, preferential occupancy of ribosomes by tRNAs at these time points correlates not only with IAV codon usage, but also with the codon usage of host antiviral response genes whose translation is occurring concomitantly. To further understand the relevance of U34 tRNA modifications in the context of IAV infection, we silenced ELP3, the catalytical unit of the Elongator complex that is responsible for the U34 modifications mcm^5^ and mcm^5^s^2^. We found that the loss of ELP3 hindered viral protein synthesis and interfered with the integrated-stress response (ISR) and unfolded protein response (UPR) related mechanisms, without inducing substantial changes in the host antiviral response. On the other hand, overexpressing ELP3 negatively affected IAV replication, likely due to a strong activation of the host immune response. Taken together, our data show that wobble uridine modifications are required for accurate IAV protein synthesis and host antiviral responses, reinforcing the relevance of the tRNA epitranscriptome in the context of viral infections.

## Results

### Host tRNA epitranscriptome is dynamic during IAV infection

To test whether IAV infection rewires the host cell tRNA epitranscriptome, adenocarcinomic human alveolar basal epithelial cells (A549) cells were infected with the influenza A/Puerto Rico/34/8 (PR8) virus strain^33^ and total RNA was isolated during a single infection cycle (from 2 to 12 hours post-infection (hpi)) **(Figure 1A)**. tRNA fractions were obtained from the extracted total RNA, and tRNA modifications were analyzed by liquid-chromatography mass spectrometry (LC-MS/MS), using a biosynthetic stable isotope labelled internal standard (SILIS), a field standard for global modification quantification^34,35^. The modification levels of 25 tRNA modifications were determined for different infection time points, namely 2, 4, 6, 8 and 12 hpi, compared to non-infected (mock) cells. Our results show that the levels of several tRNA modifications were dynamic over the infection cycle and significantly decreased at the latter stage of the viral life cycle, including ac^4^C (∼0.48 fold; p=0.0001), i^6^A (∼0.42 fold; p=0.0001), t^6^A (∼0.28 fold; p=0.0204), Q (∼0.32 fold; p=0.0057), m^2,2^G (∼0.27 fold; p=0.0010), and mcm^5^U (∼0.63 fold; p=0.0001) **(Figure 1B)**. The latter increased at early infection time-points and steadily decreased through the IAV infection cycle, while the subsequent mcm^5^s^2^U modification displayed an opposite trend **(Figure 1C)**. The introduction of mcm^5^ and mcm^5^s^2^ moieties occurs at the wobble uridine of tRNAs and is a complex enzymatic process involving different tRNA-modifying enzymes. Mechanistically, the Elongator complex (ELP1-ELP6), whose catalytic subunit relies on ELP3, initiates this enzymatic cascade by attaching a 5-carbonylmethyuridine (cm^5^U) group to the wobble uridine of eleven cytosolic tRNAs. The latter is then methylated by ALKBH8 to generate mcm^5^U bases. A subset of these mcm^5^U-modified tRNAs, namely tRNA^Lys^_UUU_, tRNA^Glu^_UUC_, tRNA^Arg^_UCU_ and tRNA^Gln^_UUG_ are further thiolated at position 2 (s^2^) to mcm^5^s^2^U by the Urm1/CTU1-2 pathway to enhance the decoding of AAA, GAA, AGA and CAA mRNA codons, respectively **(Figure 1D)**^36^. Of note, these codons are highly prevalent in the codon-biased IAV genome (**Supplementary Table 1; Supplementary Figure 1)**. The bulk tRNA LC-MS/MS data regarding mcm^5^s^2^ and mcm^5^s^2^U_34_ modifications suggest that the tRNAs carrying the mcm^5^s^2^U_34_ modification are being modified in response to IAV infection, starting early in infection with the catalysis of mcm^5^U, followed by subsequent thiolation, resulting in mcm^5^s^2^-modified wobble base. Isoacceptor-level confirmation was obtained for tRNA^Lys^_UUU_ and tRNA^Glu^_UUC_, for which we quantified and confirmed a significant increase in thiolation levels for both tRNAs at 12hpi by APM-Northern Blotting (**Figure 1E**).

**Figure 1.**
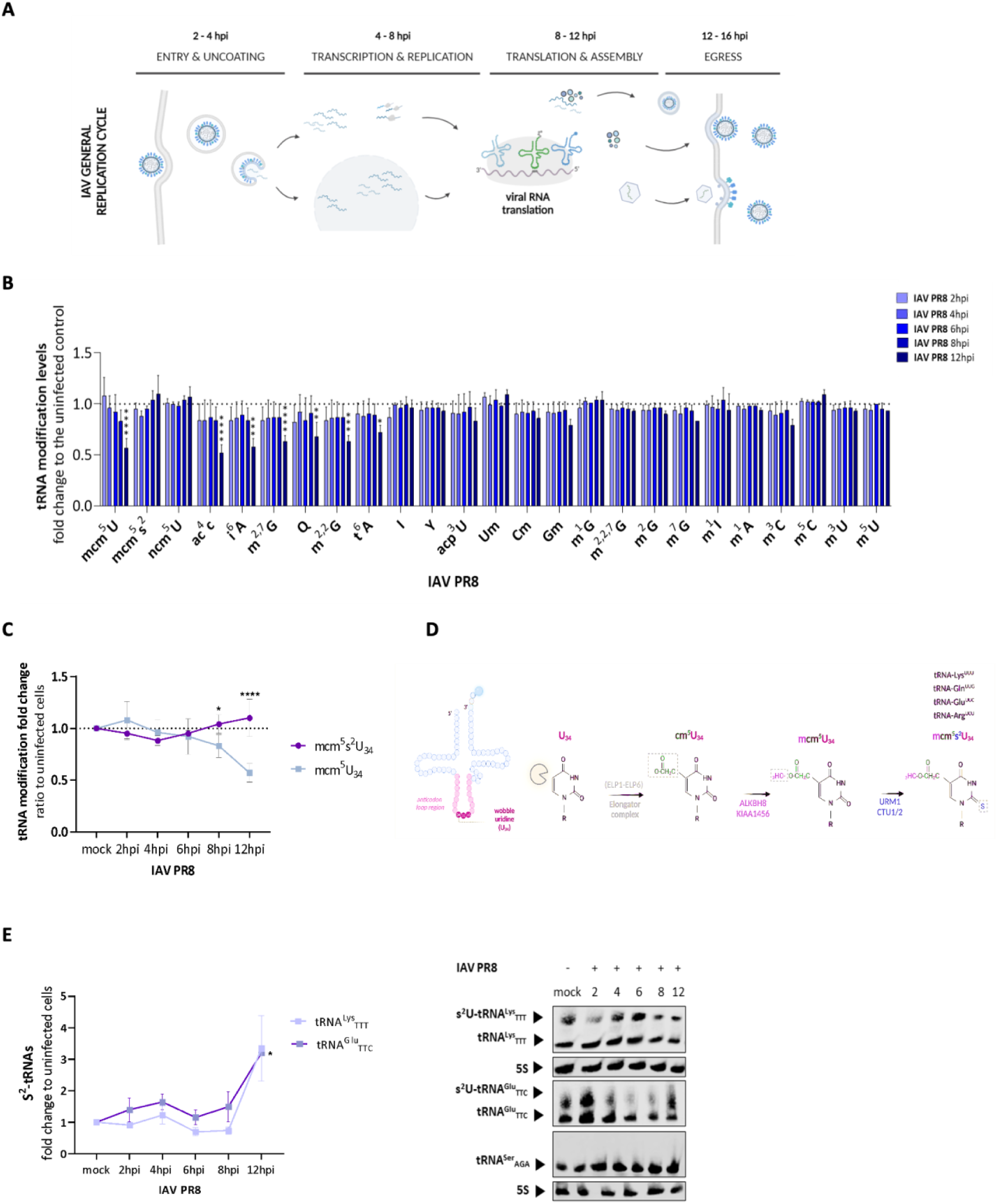
Host cell tRNA modifications and tRNA abundance profiling following IAV PR8 infection. (A) Schematic representation of the IAV life cycle, ranging between 2 to 16 hpi. (B) Quantification of tRNA modifications upon IAV PR8 infection of A549 cells by LC-MS/MS, at different infection time-points (n=3). (C) Highlight of the dynamics of mcm^5^U and mcm^5^s^2^U during IAV infection. The tRNA modification fold change in the different stages of the virus life cycle was determined in relation to mock cells (n=3). (D) Schematic representation of the enzymatic events leading to introduction of mcm^5^U and its derivatives into the wobble uridine of host cell tRNAs. (E) Quantification of s^2^ levels in tRNA^Lys^_UUU_ and tRNA^Glu^_TTC_ in mock cells and different IAV infection time-points, by APM-Northern blotting (n=2). Data analysis was conducted using one-way ANOVA, *p<0.05 and ****p<0.0001. Bars reflect the mean and error bars reflect standard deviation. A549 cells were infected at a multiplicity of infection (MOI) of 3 and samples were collected at different infection time-points, ranging between 2 to 12 hpi.

### Transcriptomic analysis identifies time-resolved host responses and dysregulation of tRNA-modification enzymes during IAV infection

To characterize host transcriptional responses to IAV infection, we performed RNA-seq of infected A549 cells at 2, 8, and 12 hpi, together with matched uninfected (mock) controls. Principal Component Analysis (PCA) revealed two distinct groups that separated mock and early-infection cells (2 hpi) from late-infection cells at 8 and 12 hpi (**Figure 2A**). This temporal separation was also reflected in the heatmap of differentially expressed genes (DEGs) (**Figure 2B**) and in volcano plots for each timepoint. We detected 311 DEGs at 2 hpi, and this number markedly increased to 3448 DEGs at 8hpi and to 4419 DEGs at 12 hpi (**Figure 2C**), indicating that substantial host transcriptional remodeling occurs only at later stages of infection. Gene Ontology (GO) enrichment analysis of DEGs at 8hpi and 12hpi highlighted pathways associated with response to stress, innate antiviral defense and host-virus interactions, consistent with robust activation of host antiviral programs (**Supplementary Figure 2A**). In contrast, early infection (2hpi) elicited minimal enrichment. Database of Clusters of Orthologous Gene (COG) function classification revealed enrichment of RNA processing and RNA modification genes at 8 and 12 hpi (**Supplementary Figure 2B**). This is consistent with altered expression of some RNA/tRNA-modifying enzymes, suggesting that IAV infection alters their expression.

**Figure 2.**
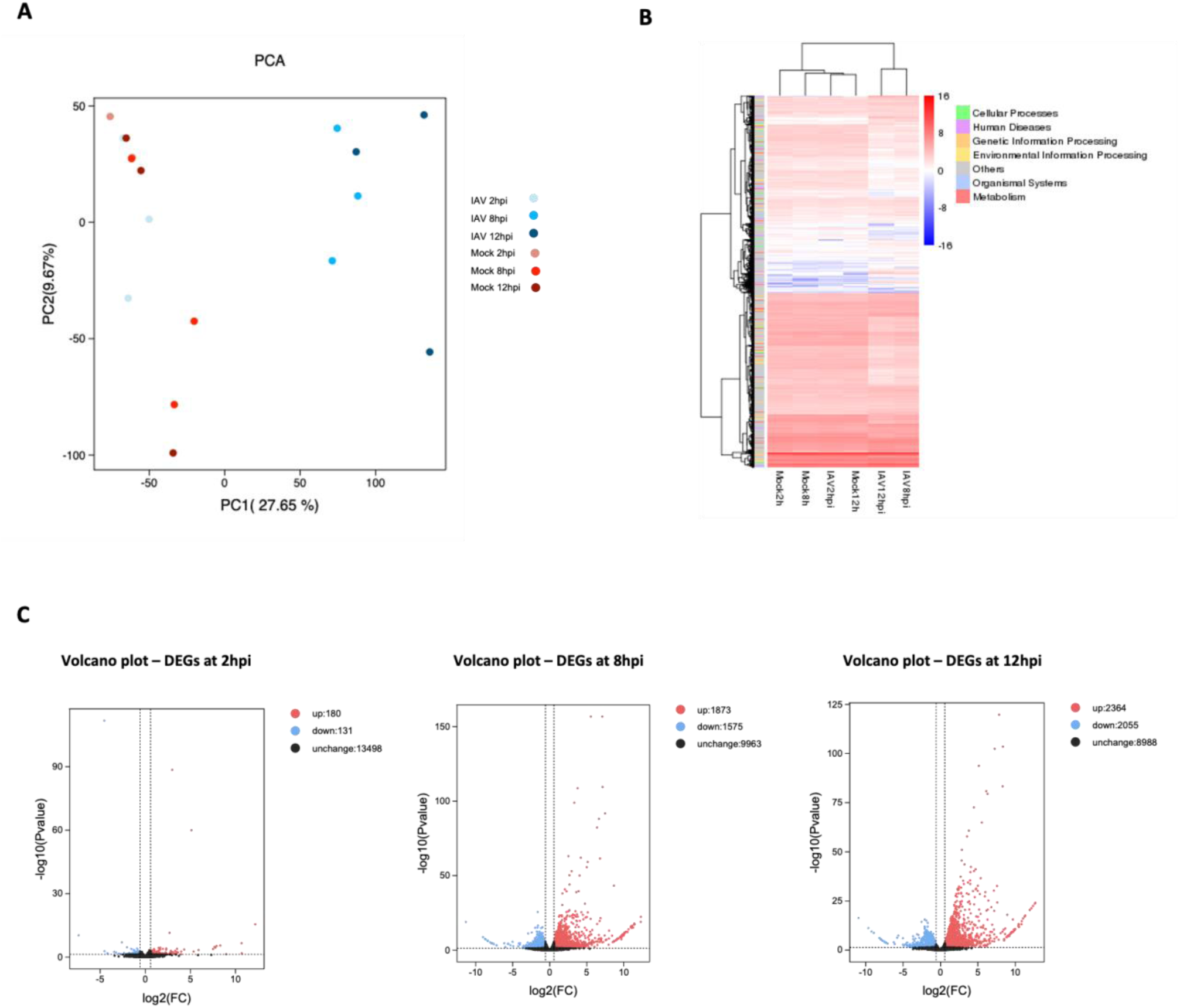
Transcriptomic differential analysis of host cells following IAV infection at 2, 8 and 12 hpi. (A) PCA analysis of all samples analyzed, including the different infection time-points and respective mocks. (B) DEGs heatmap for the different conditions analyzed (FPKM expression level normalization). A clear separation between the mock samples and 2hpi timepoints from late infection time points (8 and 12 hpi) is perceived through the analysis of both the PCA and DEG heatmap. (C) Volcano plots of DEGs at the different timepoints analyzed (2, 8 and 12 hpi). As the infection progresses, there is a steady increase in both up and down-regulated genes. Three biological replicates for each condition were sequenced, and RNA sequencing was performed using an Illumina NextSeq sequencer.

To determine whether these transcriptional changes influence codon usage patterns, we performed relative synonymous codon usage (RSCU) analysis on DEGs at each infection stage. Up-regulated genes at 8hpi exhibited a clear shift towards A-ending codons, a pattern that persisted at 12hpi (**Supplementary Figure 3**). Notably, codons such as Lys(AAA), Arg(AGA), Glu(GAA), and Gln(CAA), all decoded by mcm⁵s²U_34_ modified tRNAs and enriched in the IAV genome (**Supplementary Table 1**), were significantly overrepresented among late-stage infection up-regulated genes compared with both their synonymous counterparts (AAG, AGG, GAG, CAG). Conversely, down-regulated genes at 8 and 12 hpi showed the opposite trend, with preferential usage of AAG, AGG, GAG, and CAG, whereas AAA, AGA, GAA, and CAA were more common among genes down-regulated at 2 hpi (**Supplementary Figure 3**).

Together, these results indicate that during infection stages coinciding with active viral RNA translation (8–12hpi), the host transcriptome becomes enriched for mRNAs containing codons that are also highly represented in the IAV genome and are decoded by U34-modified tRNAs. This shift parallels the increase in mcm⁵s²U_34_ modification levels observed at late timepoints (**Figure 1C, Supplementary Figure 3**), suggesting coordinated remodeling of both host codon usage and tRNA modification pathways that favor efficient translation during IAV infection.

To further explore coordinated transcriptional programs altered during IAV infection, we performed a weighted gene co-expression network analysis (WGCNA) using the RNA-seq dataset across the analyzed infection timepoints and the non-infected controls. This analysis identified 1 co-expression module, with 118 genes, with high correlated expression patterns. This module revealed a distinct infection-stage–specific signature, showing strong positive correlation with late infection (8 and 12 hpi) and negative correlation with early infection and mock samples (**Supplementary Figure 2C**). Functional enrichment analysis demonstrated that this upregulated module at late timepoints was significantly enriched for antiviral and innate immune response pathways (**Supplementary Figure 2D**).

Further analysis of the RNA sequencing data revealed that several tRNA modifying enzymes were differentially deregulated at 8 and 12 hpi, with no alteration in expression at early infection (2 hpi). The tRNA modifying enzymes catalyzing the modifications previously identified as negatively affected by IAV infection (**Figure 1B**) were found significantly downregulated in the transcriptomic data (**Figure 3A**), namely NAT10, TRIT1, TP53RK, and Elongator complex proteins. These enzymes are involved in the catalysis of ac^4^C, i^6^A, Q, t^6^A, and cm^5^U/mcm^5^U/mcm^5^s^2^U, respectively. Surprisingly, both CTU1 and CTU2, that together act as a sulfur transferase involved in the thiolation of U34 of tRNAs, were found downregulated at both 8 and 12 hpi. On the other hand, ALKBH8, also involved in the mcm^5^ and mcm^5^s^2^U_34_ catalysis was upregulated at 12 hpi, whereas URM1, that is used by CTU1/2 complex as a sulfur donor to create the thiol group at U34 of tRNAs, was not altered upon IAV infection in any of the analyzed timepoints. Additional tRNA modifying enzymes were also found deregulated, but no effect on the corresponding tRNA modification was found (**Figure 1B**) or the corresponding tRNA modification was not analyzed.

**Figure 3.**
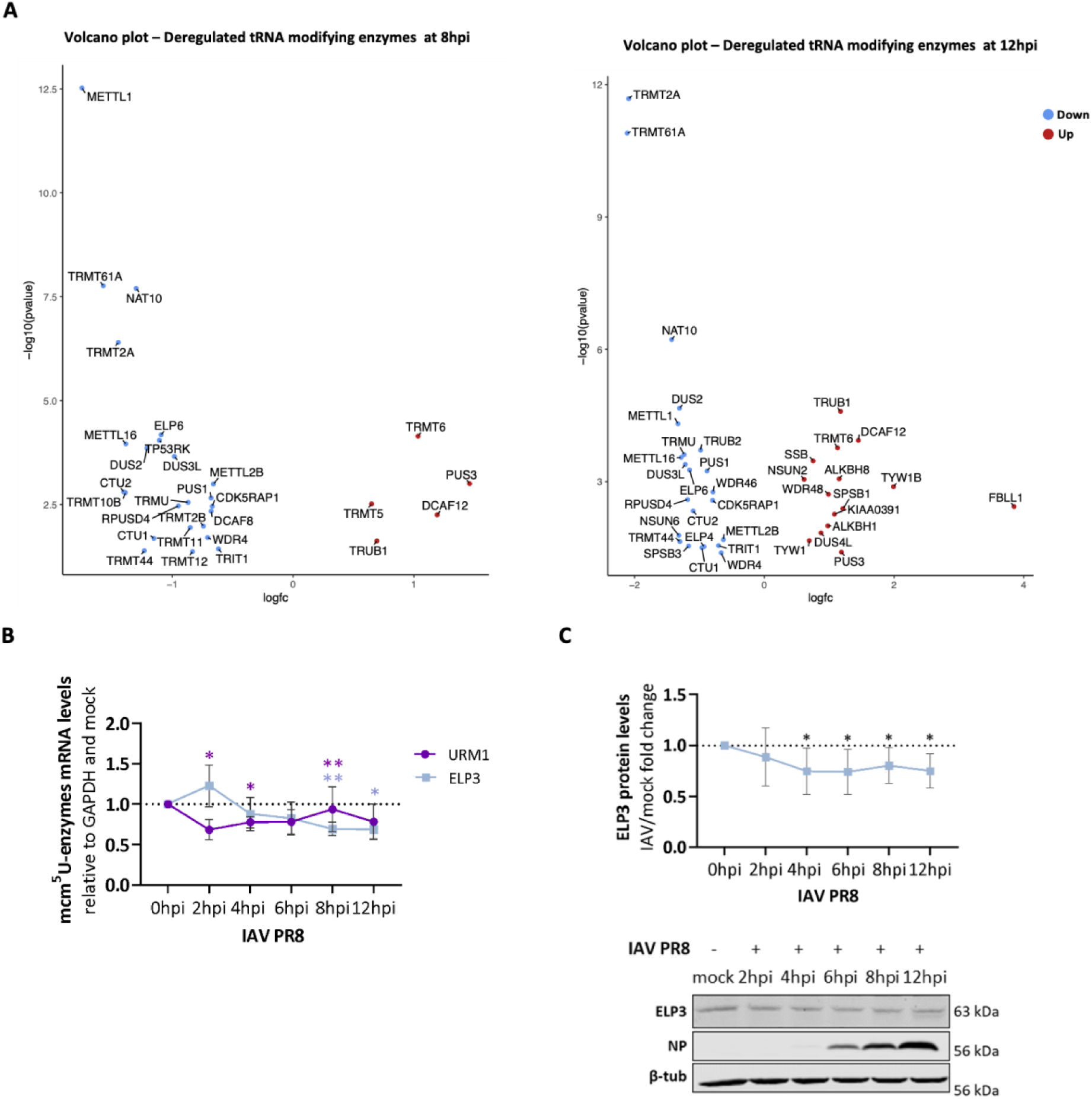
mcm^5^U and mcm^5^s^2^U catalysts are differently expressed upon IAV infection. (A) Volcano plots of the deregulated tRNA modifying enzymes found at 8 and 12 hpi in RNA-Sequencing datasets. (B) Analysis of ELP3 and URM1 abundance during the IAV PR8 infection cycle by qRT-PCR. GAPDH was used as internal control. (C) Western blot analysis of ELP3 protein levels following IAV PR8 infection. β-tubulin was used as internal control. A549 cells were infected at a MOI of 3 and samples were collected at different infection time-points, ranging between 2 to 12 hpi. Mock-infected cells were used as control. Data represents the mean of at least three independent experiments. All data analysis was performed using Student’s unpaired t-test with Welch correction, *p<0.05 and ****p<0.0001. Bars reflect the mean and error bars reflect standard deviation.

Although ELP3, the catalytical subunit of the Elongator complex, was not found as a statistically significant DEG in the transcriptomics data, we further analyzed its expression levels by both qPCR and western blotting, given its relevance for the Elongator complex, and given that other enzymes from the complex were found altered. Our data confirmed that ELP3 mRNA levels were slightly increased at early infection timepoint 2 hpi (∼0.23 fold; p=0.1763) and significantly downregulated at late infection time points, namely 8 hpi (∼0.31 fold; p=0.0051) and 12 hpi (∼0.31 fold; p=0.0135) (**Figure 3B**). Immunoblot analysis showed a statistically significant reduction of ELP3 protein levels at 4 (∼0.25 fold; p=0.0407), 6 (∼0.26 fold; p=0.0358), 8 (∼0.20 fold; p=0.0388), and 12hpi (∼0.25 fold; p=0.0147) (**Figure 3C)**, which is consistent with the steady decrease of mcm^5^U from 2 to 12 hpi **(Figure 1B)**. URM1, on the other hand, followed an opposite trend, with mRNA levels significantly downregulated at 2 hpi (∼0.32 fold; p=0.0482) and 4 hpi (∼0.2 fold; p=0.0332), but steadily recovering from 6 to 12 hpi (**Figure 3B**), in line with transcriptomics data that show no changes in URM1 expression at late infection timepoints.

### Host tRNAs that decode IAV A/U-ending codons are preferentially loaded into ribosomes during active IAV RNA translation timepoints

As altered levels of tRNA modifications could arise from changes in the abundance of specific tRNAs, we also investigated whether IAV infection perturbs global host tRNA pools using nanopore tRNA sequencing (Nano-tRNAseq)^37^. We found no specific sample clustering regardless of infection state or stage (**Figure 4A**). By quantifying the cellular tRNA levels in infected versus uninfected A549 cells, only tRNA^His^_GTG_ was found deregulated at 8hpi. No significant differences in tRNA abundances were detected in any of the other timepoints tested, indicating that the global tRNA repertoire remains stable during infection **(Figure 4C)**.

**Figure 4.**
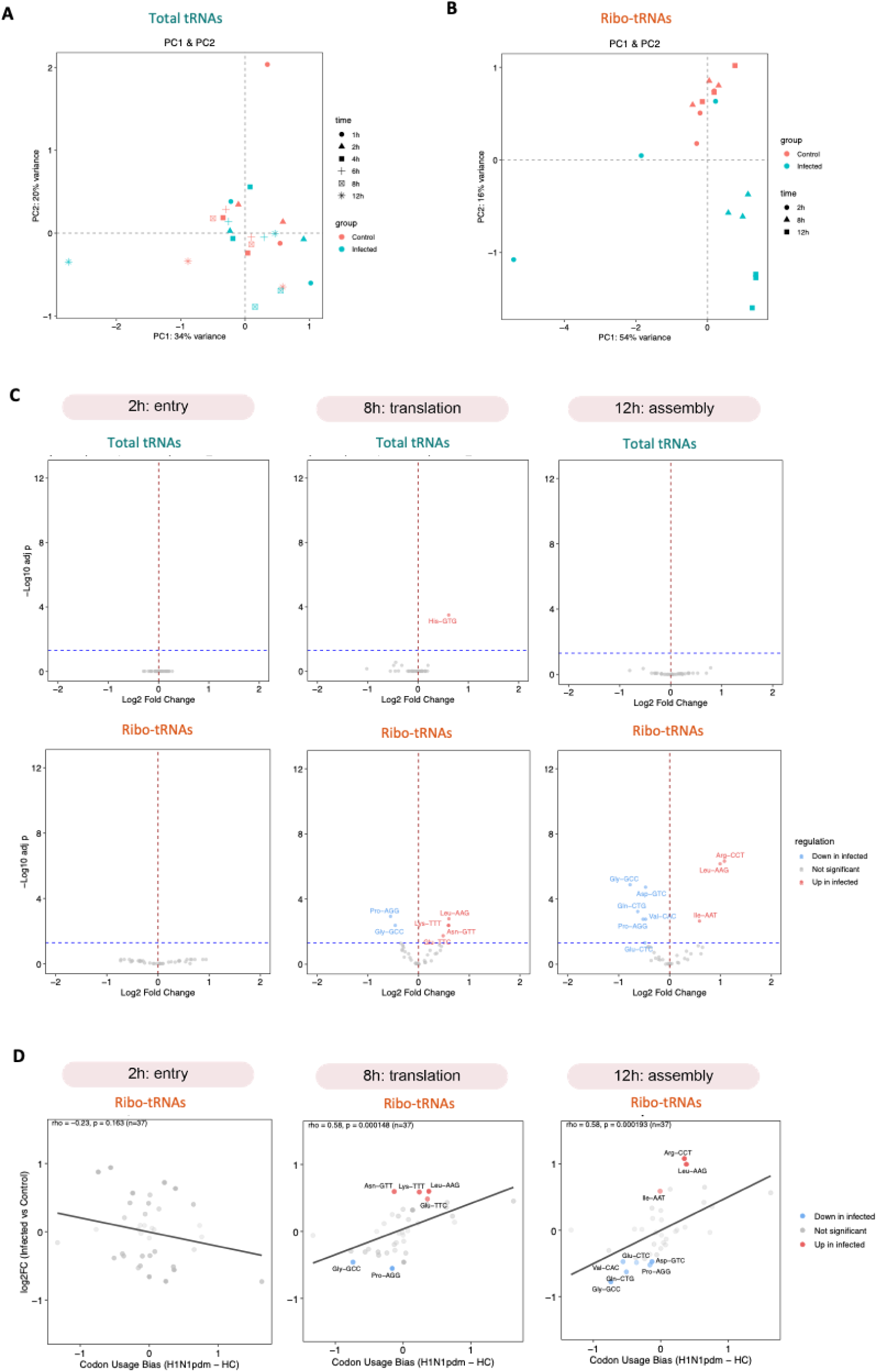
Nano-tRNASeq and Ribo-tRNASeq analysis. (A) PCA analysis of the Nano-tRNASeq experiment, corresponding to the direct tRNA sequencing of total tRNA pools in mock (control) and IAV-infected samples at different time-points (1, 2, 4, 6, 8, 12 hpi). (B) PCA analysis of the Ribo-tRNASeq experiment, corresponding to the direct tRNA sequencing of ribo-embedded tRNAs in mock (control) and IAV infected samples at different time-points (2, 8, 12 hpi). (C) Volcano plots depicting deregulated tRNAs (log2FC of infected vs mock samples) upon IAV infection at 2, 8, and 12 hpi when compared to the respective mock samples, in the total tRNA pool and in the ribo-embedded tRNA pool. Significantly upregulated tRNAs upon infection are highlighted in red, and significantly downregulated tRNAs are highlighted in blue. (D) Correlation analysis with H1N1 codon usage of ribo-embedded tRNAs in IAV infected samples at 2, 8 and 12hpi. For total tRNA analysis, 2 biological samples were analyzed by Nano-tRNASeq. For ribo-embedded tRNAs, 3 biological replicates were analyzed by Ribo-tRNASeq.

We next examined whether ribosome-associated tRNAs (tRIBO-seq^38^) would be significantly altered during active IAV RNA translation, analogous to the preferential loading of A/U-ending tRNAs upon viral infection, previously inferred from polysome-profiling studies^25^. This analysis revealed a clear shift in ribosome-bound tRNA populations (ribo-tRNAs) at 8 and 12 hpi. This temporal separation was evidenced in the PCA, which showed distinct clustering of late-infection samples (8 and 12 hpi) from mock and early-infection timepoints (2 hpi) (**Figure 4B**). Differential abundance analysis identified several ribo-tRNAs as significantly deregulated at 8 and 12 hpi, whereas no changes were detected at 2 hpi, when compared to mock samples (**Figure 4C**). Specifically, tRNA^Lys^_UUU_ and tRNA^Glu^_UUC_, whose U34 canonically carries mcm⁵s²U in human cells, were significantly enriched on translating ribosomes at 8 hpi. These tRNAs decode the highly used IAV AAA and GAA codons, respectively. On the other hand, tRNA^Gln^_CTG_ and tRNA^Glu^_CTC_, which decode CAG and GAG codons in mixed codon boxes that also contain the U34 decoded CAA and GAA codons, were significantly depleted at 12hpi **(Figure 4C**). Moreover, the ribosome-associated tRNA changes at 8 and 12 hpi correlated strongly with the codon usage bias of IAV H1N1, indicating that Ribo-tRNASeq captures virus-driven codon demand in a time-resolved manner (**Figure 4D**). Taken together, the LC-MS/MS data, the Nano-tRNASeq and the tRIBO-Seq demonstrate that tRNA populations (both in terms of modifications and abundances) are dynamically modulated in response to IAV infection, contributing to support viral protein synthesis while fine-tuning the translation machinery to meet host protein synthesis demands.

### ELP3 abundance impacts IAV mRNA translation

Since IAV transcripts are highly enriched in AAA, AGA, CAA and GAA mRNA codons, they rely heavily on tRNAs bearing Elongator-dependent modifications for efficient decoding. We therefore hypothesized that knockdown of the Elongator complex catalytic subunit ELP3 could directly impact IAV mRNA translation in a codon-dependent manner. To test this hypothesis, ELP3 was transiently silenced in A549 cells, which were then infected with IAV for 8 hours, a stage that coincides with high rates of viral RNA translation (siELP3 condition). ELP3 silencing efficiency was evaluated by qRT-PCR and western blotting and compared to the expression of ELP3 in A549 infected cells transfected with a non-targeting siRNA sequence (siCTR). The levels of ELP3 were significantly downregulated 48 hours post-transfection both at the mRNA (∼0.9 fold; p=0.0038) and protein levels (∼0.9 fold; p=0.0025), confirming the efficiency of the knockdown **(Figure 5A)**. Of note, no major alterations on cellular viability were observed **(Supplementary Figure 4A)**. To confirm that the loss of ELP3 induces tRNA hypomodifications, the levels of tRNA modifications were quantified by LC-MS/MS^35,39^. As anticipated, the ELP3 knockdown induced a significant decrease in the levels of mcm^5^U (∼0.5 fold; p<0.0001) and its derivative mcm^5^s^2^U (∼0.3 fold; p<0.0001) when compared to the siCTR condition **(Figure 5B)**. No significant alterations in the remaining quantified tRNA modifications were observed in both infection and non-infection contexts **(Supplementary Figure 4B)**, highlighting the relevance of ELP3 in writing these wobble modifications. To rule out the possibility of the tRNA abundance itself impacting thiolation levels found per tRNA upon ELP3 silencing, we quantified the total levels of tRNA^Glu^_UUC_, tRNA^Gln^_UUG_ and tRNA^Lys^_UUU_ isoacceptors by Northern blotting and, regardless of IAV PR8 infection, no major alterations in tRNA levels were observed in the absence of ELP3 **(Figure 5C)**.

**Figure 5.**
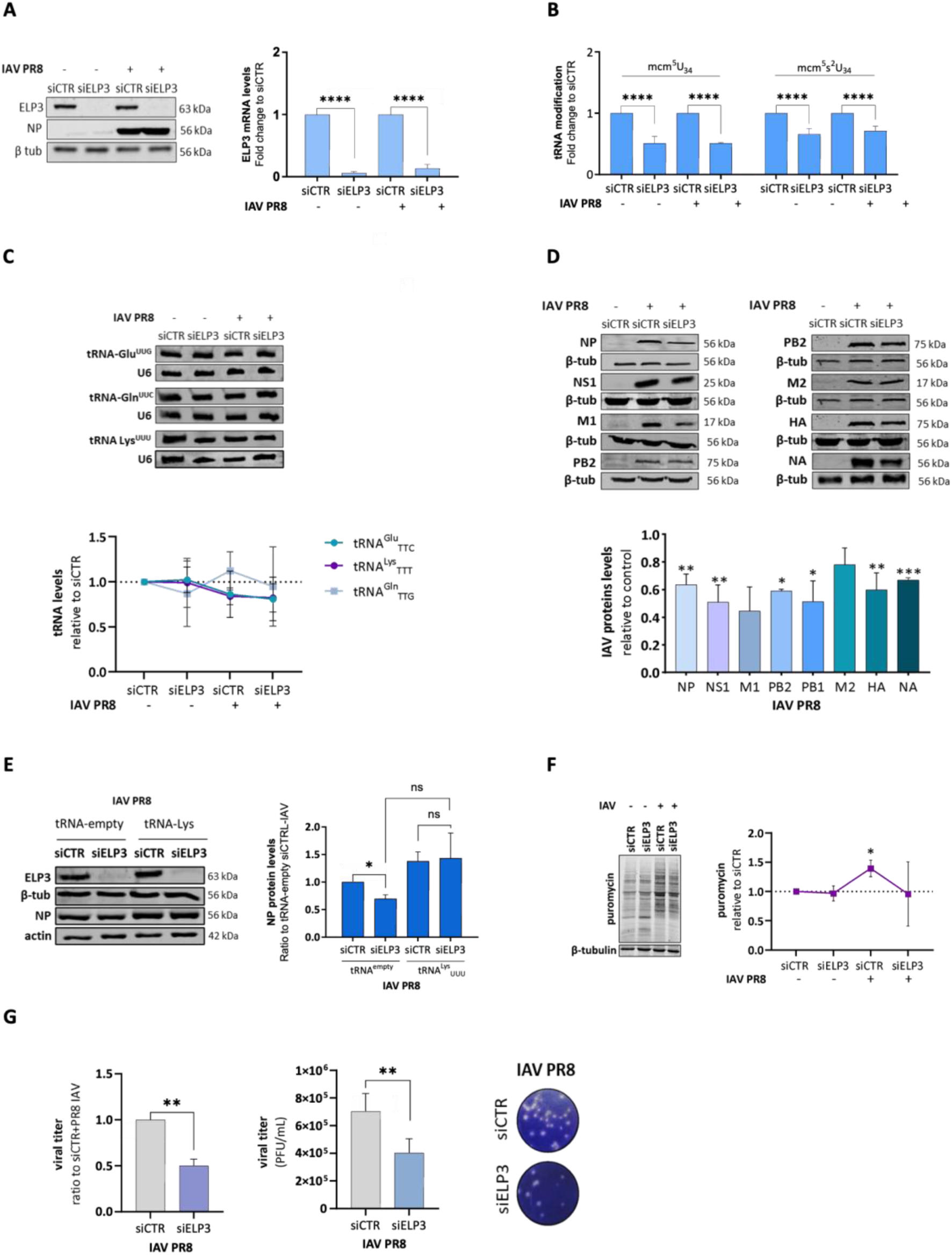
Loss of ELP3 induces tRNA hypomodifications and hinders IAV genome translation. (A) Assessment of ELP3 silencing efficiency by RT-qPCR and Western Blotting. (B) Quantification of tRNA modifications upon the ELP3 knockdown by LC- MS/MS. (C) Assessment of total tRNA^Lys^_UUU_, tRNA^Gln^_UUG_, and tRNA^Glu^ levels by Northen blotting. U6 rRNA was used as internal control. (D) Impact of the ELP3 silencing in viral protein synthesis by western blotting. (E) Effect of the tRNA^Lys^_UUU_ overexpression in ELP3-depleted cells following IAV PR8 infection. (F) Effect of the ELP3 knockdown in cellular protein synthesis by western blotting. (G) Quantification of viral titers following IAV PR8 infection of ELP3 knocked-down cells by plaque assay. A549 cells were infected at a MOI of 3 and all samples were collected at 8 hpi. For plaque assays, IAV-infected cells were collected at 16 hpi. Mock-infected cells were used as control. Data represents the mean of at least three independent experiments. Bars reflect the mean and error bars reflect standard deviation. All data analysis was performed using Student’s unpaired t-test with Welch correction, *p<0.05 and ****p<0.0001.

To evaluate the effect of the ELP3 knockdown in IAV genome decoding we measured the levels of several IAV proteins by immunoblotting. As shown in **Figure 5D**, the inhibition of ELP3 led to a significant decrease in the abundance of different viral proteins, including NP (∼0.4 fold decrease; p=0.0027), HA (∼0.4 fold decrease; p=0.0077), NA (∼0.3 fold decrease; p=0.0009), PB1 (∼0.4 fold decrease; p=0.0303), PB2 (∼0.4 fold decrease; p=0.0155), and NS1 (∼0.5 fold; p=0.0042). To confirm that the reduced viral protein production induced by the ELP3 deficiency was due to mcm^5^U and mcm^5^s^2^U hypomodifications, we transfected siCTR- and siELP3-treated cells with a tRNA^Lys^_UUU_ plasmid. If the levels of viral codon-biased mRNA translation would be restored following overexpression of one of the mcm^5^s^2^U-modified tRNAs, this would demonstrate the central role of these modifications in the recognition of IAV mRNA codons, as increasing tRNA abundance compensates for the absence of wobble uridine modifications^9,15,40^. Regardless of prior ELP3 silencing, the tRNA^Lys^_UUU_ plasmid was effectively transfected in A549 cells **(Supplementary Figure 4C)**. As shown in **Figure 5E**, transfection of siCTR-treated cells with the tRNA^Lys^_UUU_ increased the protein levels of NP, although not statistically significant (∼0.4 fold; p=0.0619), demonstrating that increasing the levels of this tRNA in cells boosts the recognition of its cognate mRNA codon, enriched in IAV genes. Importantly, the protein levels of NP were recovered by overexpression of the tRNA^Lys^_UUU_ plasmid in ELP3-depleted cells compared to the counterpart overexpressing an empty-tRNA plasmid **(Figure 5E)**.

To determine the extent of which the ELP3 knockdown impacts host cell protein synthesis, we also monitored and quantified nascent protein synthesis rates using the puromycin incorporation SUnSET method^41^. No statistically significant differences in cellular protein synthesis rates were observed in siELP3- over siCTR-treated cells **(Figure 5F)**. However, IAV PR8 infection of A549 cells induced a ∼1.6 fold (p=0.0276) increase in cellular protein synthesis (**Figure 5F**). This effect was partially counteracted by the ELP3 knockdown following IAV PR8 infection **(Figure 5F**). This protein synthesis reduction is similar to the one observed in ELP3 silenced cells when compared to non-silenced cells, in the absence of IAV infection. Of note, the protein levels of several common non-codon biased (C/G-ending codons) host housekeeping genes were not significantly affected by the inhibition of ELP3, either in the infectious or non-infectious context **(Supplementary Figure 4D)**. Collectively, these results support that mcm^5^U and mcm^5^s^2^U hypomodifications affect the decoding of biased A-ending IAV’s mRNAs.

To further verify if depletion of ELP3 affects the propagation of IAV, we quantified the yields of infectious virus particles using plaque assays. Our results demonstrated that there was a statistically significant reduction (∼0.5 log; p=0.0069) in the number of produced infectious viral particles **(Figure 5G)**, indicating that the ELP3 knockdown perturbs the IAV propagation.

Although A549 cells are commonly used to study IAV infection, we also considered the use of another epithelial lung cell line derived from, non-tumorigenic lung tissue to exclude possible side effects at the epitranscriptome level. We infected BEAS-2B cells with IAV PR8 and analyzed the abundance of ELP3 within a single cycle infection up to 8hpi. When comparing uninfected BEAS-2B with BEAS-2B IAV PR8-infected cells, we found that the ELP3 mRNA levels were significantly up-regulated at 4hpi (∼0.5 fold; p=0.0420) (**Supplementary Figure 5A**), instead of at 2hpi, as observed for A549 IAV infected cells (**Figure 3B**). ELP3 expression levels decreased steadily up to 8 hpi to near basal levels in BEAS-2B (**Supplementary figure 5A**). Of note, basal ELP3 levels in BEAS-2B cells are lower than in A549, which may explain the slight differences of ELP3 expression throughout one IAV infection cycle (**Supplementary Figure 5B**). Despite this, silencing of ELP3 prior to IAV PR8 infection of BEAS-2B cells significantly reduced the protein levels of NP (∼0.4 fold; p=0.0374) (**Supplementary Figure 5C),** and decreased the viral titer to half (**Supplementary Figure 5D**) thus recapitulating our findings in A549 cells.

### ELP3 levels differentially modulate host immune response pathways

The host cell has developed multiple strategies to counteract IAV infection^42,43^. A central component of this response is the RIG-I/MAVS pathway, which detects cytoplasmic viral RNA and triggers the induction of interferons (IFNs) and numerous interferon-stimulated genes (ISGs), including IRF1^44,45^. To infer if the RIG-I/MAVS signaling was affected by the inhibition of ELP3, siCTR- and siELP3-treated cells were transfected with the RIG-I agonist 3p-hairpin RNA (3p-hpRNA) for 6 hours, as previously described^46^. Following stimulation, our data shows that IFN-β and IRF1 mRNA levels were robustly induced relative to unstimulated cells, but no significant differences were detected between siCTR and siELP3 conditions (IFN-β: p=0.5506; IRF1: p=0.7934) (**Figure 6A**). These findings indicate that ELP3 knockdown does not impair RIG-I/MAVS pathway activation. In contrast, when ELP3-silenced cells were infected with IAV PR8, we observed a modest but significant up-regulation of IFN-β (∼2 fold; p=0.0275). Although IRF1 mRNA and protein levels were consistently upregulated across all infected conditions relative to mock controls, no differences were observed after IAV infection in ELP3 silenced cells when compared to siCTR-infected cells (**Figure 6B, C**). Thus, ELP3 depletion does not amplify the host cell antiviral response through the RIG-I/MAVS pathway, and the reduction in viral titers in siELP3-silenced cells is unlikely to result from enhanced innate immune activation. Additionally, STAT1 activation assessed by its phosphorylation status, remained unchanged in siELP3-infected cells compared with siCTR-infected cells, and IFIT1 was significantly downregulated **(Figure 6D)**. This supports the conclusion that ELP3 knockdown does not activate canonical antiviral signaling pathways.

**Figure 6.**
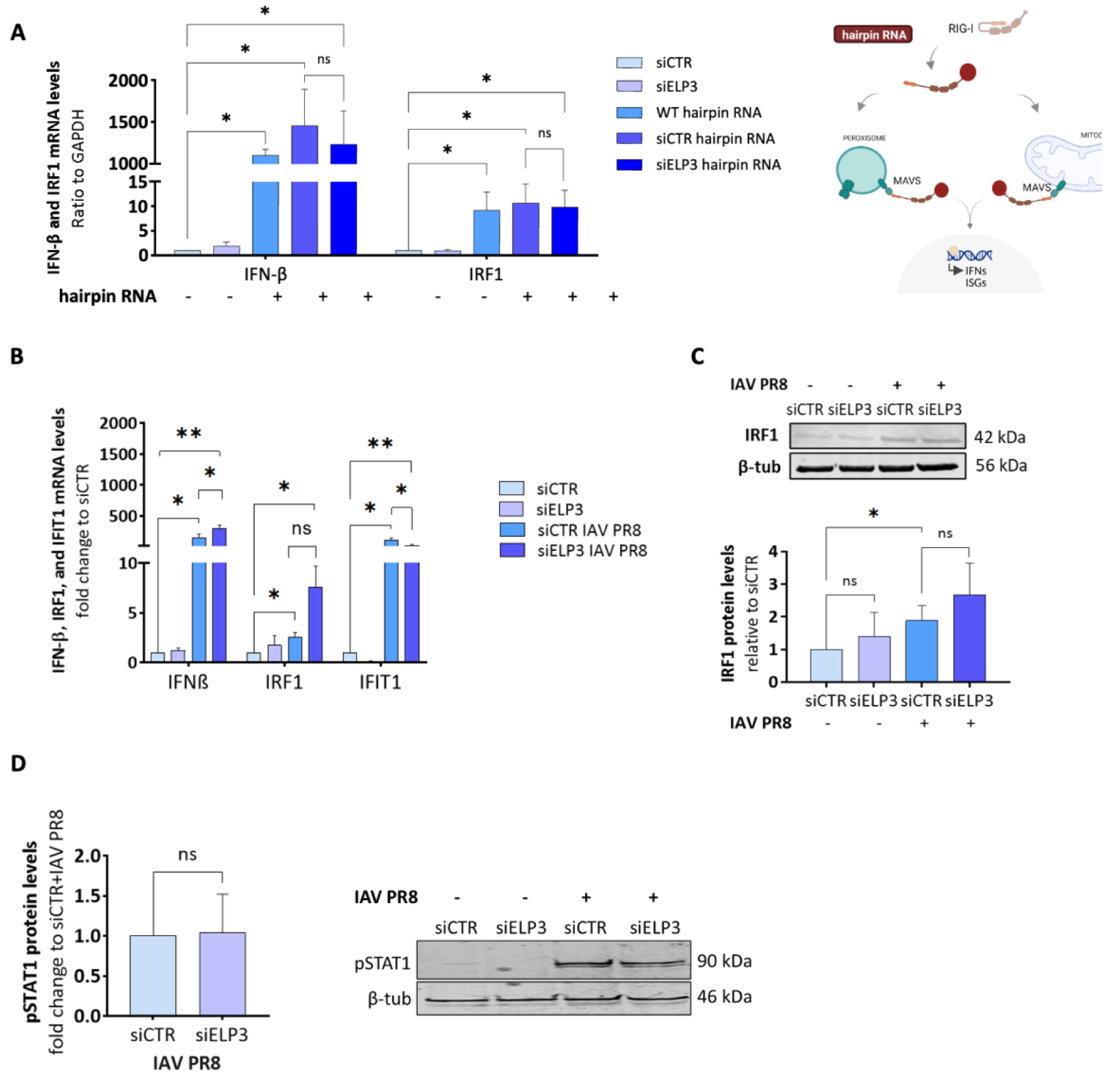
Effect of ELP3 loss in host antiviral responses. (A) Effect of the ELP3 knockdown on the RIG-I/MAVS signaling pathway following its activation with a 3p-hairpin RNA molecule. (B) Analysis of IFN-β, IRF1 and IFIT1 mRNA levels following IAV PR8 infection of ELP3-depleted cells. (C) Western blot analysis of IRF1 abundance following IAV infection. (D) Effect of the ELP3 knockdown on the phosphorylation status of STAT1. A549 cells were infected at a MOI of 3 and collected at 8 hpi. Data represents the mean of at least three independent experiments. Mock-infected cells were used as control. Bars reflect the mean and error bars reflect standard deviation. All data analysis was performed using Student’s unpaired t-test with Welch correction, *p<0.05 and ****p<0.0001.

Since our transcriptomics analysis revealed a shift toward A-ending codons in up-regulated host genes at late infection timepoints and enrichment of immune- and antiviral-related pathways, we next investigated whether ELP3 overexpression influences host defense responses (**Supplementary Figure 6A**). Remarkably, overexpression of ELP3 in A549 cells prior to IAV infection led to a significant increase in IRF-1, IFN-β and IFIT1 (**Supplementary Figure 6B**), and a significant increase in phosphorylated STAT1 (p-STAT1), accompanied by reduced viral titers and lower IAV NP protein levels (**Supplementary Figure 6C**), when compared to the control transfected with an empty plasmid. Notably, ELP3 overexpression alone, even in the absence of infection, was sufficient to induce p-STAT1 (**Supplementary Figure 6D**). This suggests that ELP3 elevation can prime or potentiate antiviral signaling pathways, thereby contributing to reduced viral replication. Taken together, our data suggest that a physiological range of U34 modification supported by ELP3 is required for accurate IAV protein synthesis, and that outside this range, elevated ELP3 can prime antiviral signaling and suppress replication.

### Loss of ELP3 is associated with ISR hyperactivation and attenuates the IAV-induced activation of the UPR IRE1α/XBP1 axis

Decreased levels of ELP3 are associated with activation of the ISR and the UPR due to tRNA hypomodifications^8,15,47^, and several studies have also proposed a link between ER stress responses and innate immunity as reviewed before^48^. We have recently found that, to facilitate its propagation, IAV induces the accumulation of insoluble proteins and ER stress responses via the UPR-IRE1α-XBP1 pathway, with increased phosphorylation of eIF2α^46^.

We questioned the extent to which the loss of ELP3 could affect the ISR- and UPR- related mechanisms during IAV infection, crucial for the virus propagation^46,49^. A simplified view of these mechanisms is depicted in **Figure 7A**. We found that the ELP3 knockdown induced the phosphorylation of eIF2α (∼0.4 fold; p=0.0362), along with an increased GADD34 expression (∼2.6 fold; p=0.0713) in the non-infected conditions **(Figure 7B; top panel)**. IAV PR8 infection also induced phosphorylation of eIF2α (∼2 fold; p=0.0052) and further induced the phosphorylation of ATF4 (∼2 fold; p=0.0056). As expected, infection of ELP3-depleted cells had a cumulative effect, exacerbating the phosphorylation of eIF2α (∼3 fold; p=0.0362) and ATF4 (∼3 fold; p=0.0490). However, for GADD34, this cumulative effect was not observed in comparison to infected cells that were not previously ELP3 silenced **(Figure 7B; top panel).** To assess if the IRE1α-XBP1 pathway was also affected by the inhibition of ELP3, we monitored the ratio of unphosphorylated and phosphorylated IRE1α by immunoblotting and the mRNA levels of sXBP1 by qRT-PCR. We confirmed that this UPR-branch was activated following IAV PR8 infection, as demonstrated by the increased phosphorylation of IRE1α (∼2 fold; p=0.0290) and splicing of XBP1 (∼4.5 fold; p=0.0101) at 8 hpi **(Figure 7B; bottom panel)**. Remarkably, the levels of IRE1α phosphorylation (∼0.7 fold; p=0.0311) and the mRNA levels of sXBP1 (∼0.4 fold; p=0.0494) were significantly decreased following IAV PR8 infection of ELP3-knocked cells in comparison with siCTR-treated cells **(Figure 7B; bottom panel)**. Notably, overexpression of a tRNA^Lys^_UUU_ in IAV-infected and ELP3-depleted cells was sufficient to rescue eIF2α phosphorylation and XBP1 splicing near to IAV-induced basal levels, demonstrating that the ELP3 knockdown disrupts proteostasis due to tRNA hypomodifications (**Figure 7C,D)**.

**Figure 7.**
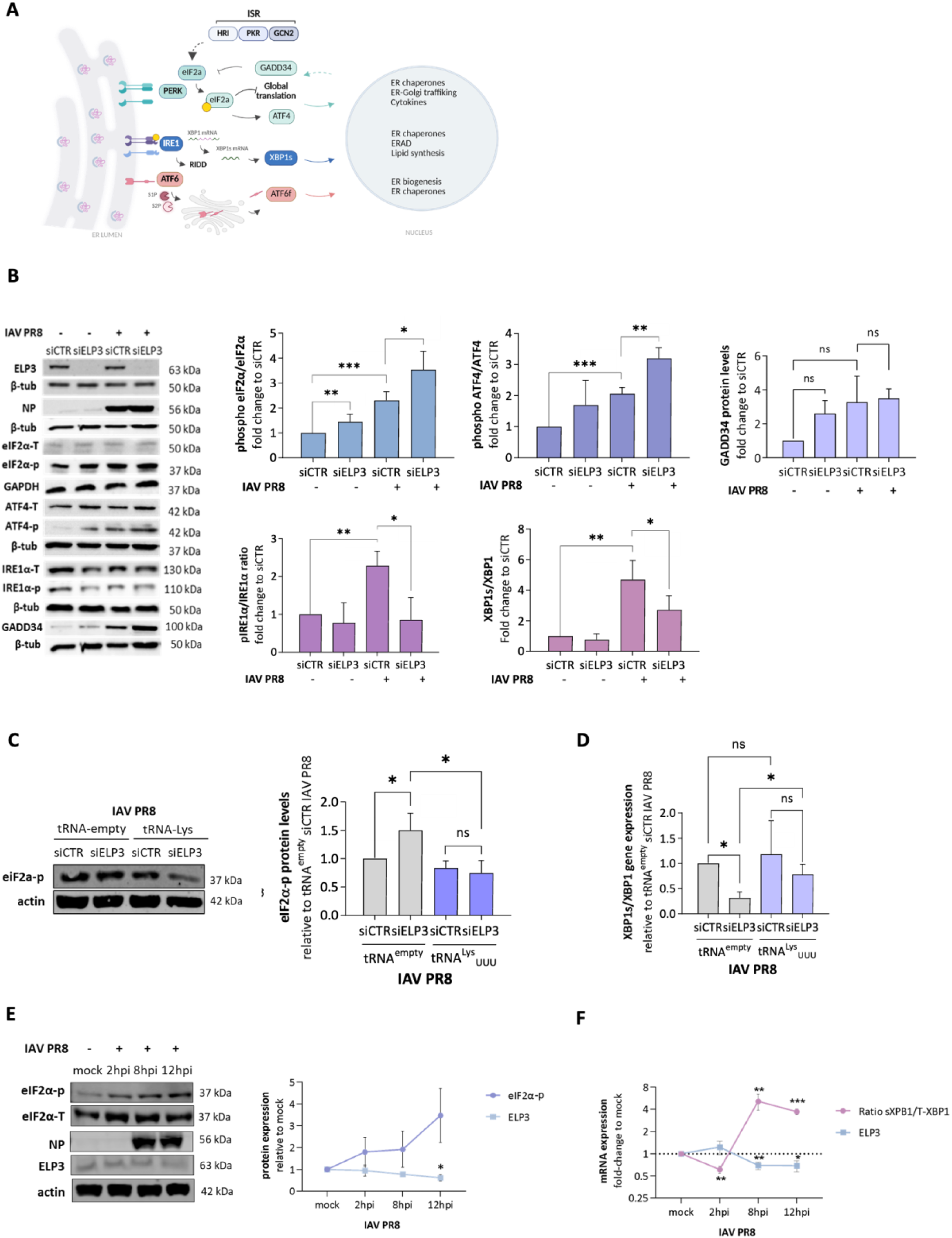
ELP3 inhibition activates the ISR and interferes with UPR-related mechanisms essential to IAV propagation. (A) Simplified view of the UPR- and ISR-related mechanisms. (B) Top panel: Effect of the knockdown of ELP3 on the ISR by immunoblotting. Bottom panel: IRE1α-UPR branch by immunoblotting and qRT-PCR. (C) Immunoblot analysis of eIF2α phosphorylation levels following tRNA^Lys^_UUU_ overexpression in tRNA hypomodification conditions. (D) qRT-PCR analysis of sXBP1 levels following tRNA^Lys^_UUU_ overexpression in tRNA hypomodification conditions. (E) Analysis of eIF2α phosphorylation and ELP3 protein levels at different stages of IAV PR8 infection by immunoblotting. (F) Analysis of sXBP1 and ELP3 mRNA abundance at different stages of IAV PR8 infection. A549 cells were infected at a MOI of 3 and samples were collected at 8hpi. Data represents the mean of at least three independent experiments. Bars reflect the mean and error bars reflect standard deviation. All data analysis was done using Student’s unpaired t-test with Welsch correction, *p<0.05 and ****p<0.0001.

We further monitored the levels of eIF2α and IRE1α at specific time-points within IAV infection (2, 8, and 12 hpi) and evaluated their correlation with the abundance of ELP3. In agreement to our previous results^46^, we found that phosphorylation of eIF2α steadily increases until 12 hpi **(Figure 7E)**. On the other hand, while at early infection time points the expression of sXBP1 is significantly reduced (∼0.4 fold; p=0.0013), the levels of sXPB1 are significantly increased at 8 hpi (∼5 fold; p=0.0070) when compared to the uninfected control. At 12 hpi, although the levels of sXBP1 remained elevated (∼3 fold; p=0.0005), they were lower than at the 8 hpi time point **(Figure 7F)**. The expression of ELP3 changed in line with these observations, being significantly downregulated at 12 hpi (∼0.4 fold; p=0.0433), coinciding with the peak of p-eIF2α **(Figure 7E,F)**. These results confirm that the abundance of ELP3 is mostly affected during the latter stages of IAV infection, in which the virus is replicating at higher levels of ER stress compared to the early stages of infection. These observations indicate that the reduced abundance of ELP3 during the infectious cycle could be a consequence of the IAV-induced ER stress. To clarify this, we challenged A549 cells with two different stress-inducing agents, including sodium arsenite (SA) and dithiothreitol (DTT), between 0.3 to 3 hours. We further investigated how these stressors affected ER stress responses and how this correlated with the abundance of ELP3. Under our experimental settings, exposure of A549 cells to DTT led to eIF2α phosphorylation at 0.3 hours post treatment (hpt) (∼2 fold; p=0.0504). This response persisted throughout the entire treatment. GADD34 expression (∼4 fold; p=0.1272) exceeded the levels of eIF2α phosphorylation at 3 hpt **(Supplementary Figure 7A top panel)**. Moreover, the protein levels of ELP3 were slightly decreased in the onset of treatment, at 0.3 (∼0.15 fold; p=0.0204) and 1 (∼0.25 fold; p=0.3086) hpt, returning to baseline levels at 3 hpt (p=0.8995) **(Supplementary Figure 7A top panel)**. As expected, following DTT treatment, decreased expression of ELP3 was in line with the induction of eIF2α phosphorylation (∼0.4-fold increase; p=0.1690) **(Supplementary Figure 7B)** and, though not statistically significant, inhibited the splicing of XBP1 (∼2.8 fold; p=0.2514) when compared to DTT-siCTR-treated cells (∼3.5 fold induction; p=0.2514) **(Supplementary Figure 7C)**.

Similarly to DTT, incubation with SA induced eIF2α phosphorylation after 30 minutes (∼1.5 fold; p= 0.3956) and 1 hour (∼2 fold; p=0.0080) treatment. Though still elevated, the levels of eIF2α phosphorylation began to decline after 2 hours of treatment, coinciding with increased expression of GADD34 (∼3.5 fold; p=0.1018) **(Supplementary Figure 7A, bottom panel)**. ELP3 abundance mirrored the pattern observed for DTT treatment, being reduced after 30 minutes (∼0.5 fold; p=0.0987) and 1-hour (∼0.6 fold; p=0.0345), starting to recover at 2hpt (∼0.35 fold; p=0.1036) and 3hpt (∼0.3 fold; p=0.3187) **(Supplementary Figure 7A, bottom panel)**. Loss of ELP3 was followed by an increased phosphorylation of eIF2α after SA treatment over SA-treated-siCTR cells (∼0.1 fold increase; p=0.3059) **(Supplementary Figure 7B)**. Splicing of XBP1 was not induced after SA treatment, being reduced in both siCTR (∼0.6 fold; p=0.1131) and siELP3 (∼0.6 fold; p=0.1468) conditions **(Supplementary Figure 7C)**. Collectively, these results demonstrate that ELP3 functions as a critical integrator of tRNA epitranscriptomic status and ER stress responses, and that its loss diverts the host response toward ISR activation while preventing full engagement of the pro-viral IRE1α–XBP1 axis during IAV infection.

### ELP3 favors the propagation of another codon biased RNA virus, the vesicular stomatitis virus

To assess if other viruses exploit ELP3 and its dependent modifications to the same extent as IAV, we infected A549 cells with the non-segmented negative-strand RNA vesicular stomatitis virus (VSV), another virus with a bias to AAA, AAG, and AAC mRNA codons **(Figure 8A)**. We analyzed the dynamics of ELP3 abundance at various time-points within a single VSV infection cycle and found that, at the mRNA level, ELP3 expression was significantly decreased at 12 hpi (∼0.3 fold; p=0.0314), when compared to the uninfected control **(Figure 8B, left panel)**. On the other hand, ALKBH8 mRNA expression was already significantly decreased at 4 hpi (∼0.26 fold; p=0.0238), and at 12 hpi (∼0.67 fold; p=0.0094) **(Figure 8B, left panel).** URM1 expression was consistently downregulated at 6 (∼0.6 fold; p=0.0039), 8 (∼0.6 fold; p=0.0556), and 12hpi (0.62 fold; p= 0.0088) **(Figure 8B, left panel)**.

**Figure 8.**
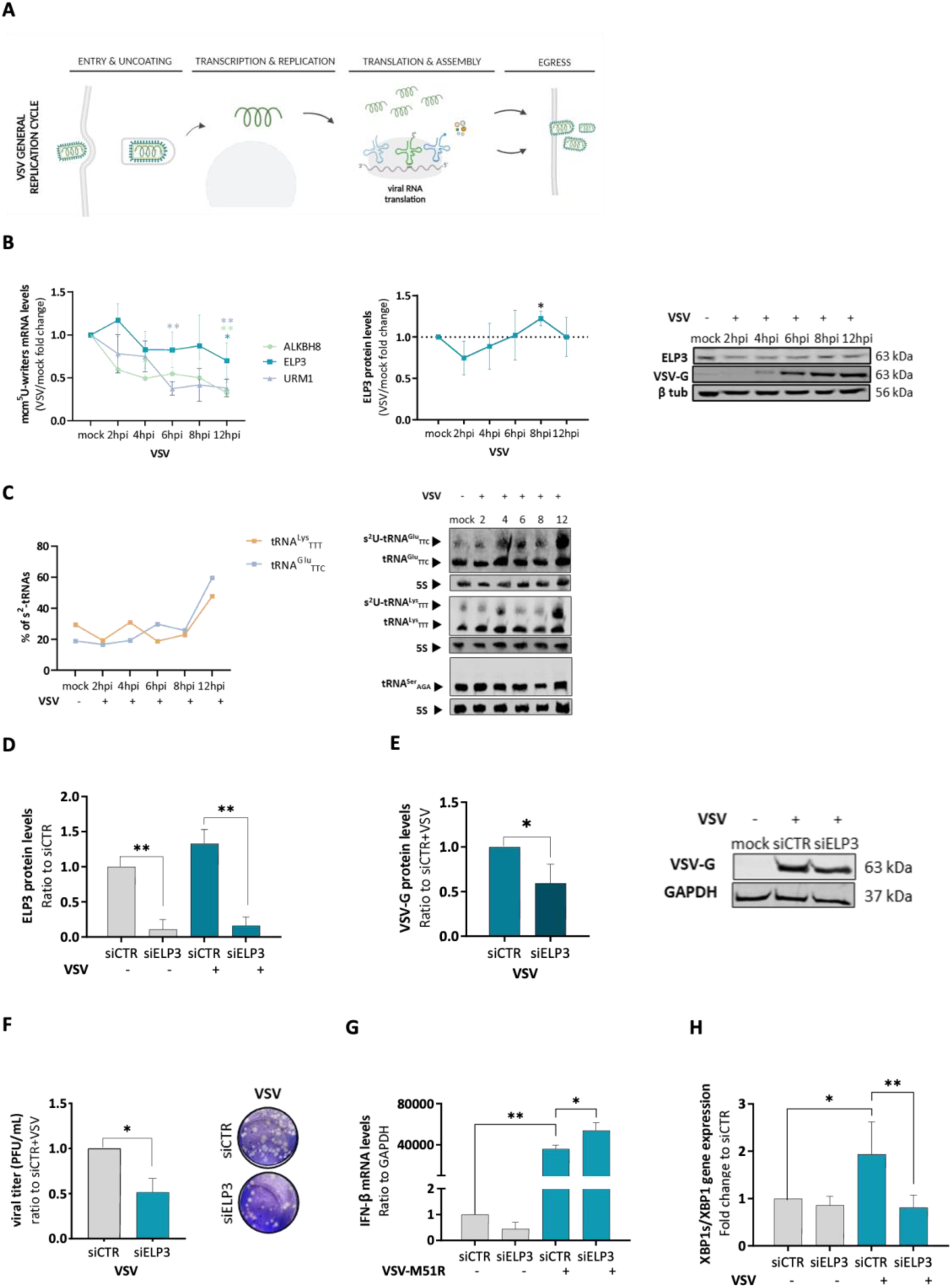
Loss of ELP3 affects the decoding and propagation of VSV. (A) Schematic representation of the VSV infectious cycle. (B) tRNA-modifying enzyme abundance within a single VSV infection cycle by qRT-PCR. (C) Abundance of mcm^5^s^2^U_34_ in specific tRNA isoacceptors over the course of VSV infection by northern blot. (D) ELP3 silencing efficiency by western blotting. (E) Effect of the ELP3 knockdown in the protein levels of VSV-G by Western blotting. (F) Effect of the ELP3 knockdown in VSV infectious viral particle production by the plaque assay. (G) Effect of the ELP3 knockdown in the mRNA levels of IFN -β after VSV-M51R infection by qRT-PCR. VSV-M51R carries the M51R mutation in the VSV matrix protein, which is responsible for inhibiting several host pathways related to cytokine signaling and IFN responses. (H) Effect of the ELP3 knockdown in the induction of sXBP1 upon VSV infection. A549 cells were infected with VSV or VSV-M51R with a MOI of 3. For plaque assays, samples were analyzed at 16hpi. For the remaining experiments, cells were collected at 8hpi. Data represents the mean of at least three independent experiments. Bars reflect the mean and error bars reflect standard deviation. All data analysis was done using Student’s unpaired t-test with Welch correction, *p<0.05 and ****p<0.0001.

To quantify the thiolation levels in tRNAs, we performed N-acryloylamino phenyl mercuric chloride (APM) gel electrophoresis combined with Northen blotting^50^ for tRNA^Glu^_UUC_ and tRNA^Lys^_UUU_ upon VSV infection **(Figure 8C)**. As a negative control, we analyzed the thiolation levels of tRNA^Ser^_AGA_ as it is not thiolated and hence it is not modified with mcm^5^s^2^U. We found that the levels of thiolated tRNA^Glu^_UUC_ were significantly higher at 12 (∼4.5 fold increase; p=0.0003) hpi **(Figure 8C).** The levels of thiolated tRNA^Lys^_UUU_ were also increased at 12 hpi, although not statistically significant, compared to the non-infected control **(Figure 8C)**. This indicates that the reduced URM1 mRNA levels do not affect the modification catalysis during VSV infection, and other enzymes such as CTU1 or CTU2 may be involved.

To delve deeper into the function of ELP3 during VSV genome decoding, we knocked-down the expression of ELP3 using siRNAs **(Figure 8D)** and observed a marked reduction in the levels of the viral VSV-G protein (∼0.4 fold; p=0.0321) as anticipated **(Figure 8E)**, indicating that ELP3 and its dependent tRNA modifications are also relevant to an effective decoding of VSV. The silencing of ELP3 also affected VSV titers as shown by the decrease (50%; p=0.013) in the amount of VSV infectious viral particles **(Figure 8F)**. Following infection of ELP3-silenced cells with VSV, the mRNA levels of IFN-β were also significantly increased (∼0.5 fold; p=0.0360) over those of siCTR-treated cells **(Figure 8G).** Similarly to IAV infection, VSV infection induced the UPR-IRE1α branch, as depicted in **figure 8H** by the increased splicing of XBP1 (∼0.94 fold; p=0.0196). As expected, the knockdown of ELP3 blocked the induction of sXBP1 upon VSV infection (∼0.4 fold; p=0.0080) **(Figure 8H)**. These results suggest that this UPR-related mechanism is also activated in the context of VSV infection and is affected by the suppression of ELP3 expression.

## Discussion

Different reports have identified tRNA modifications as virulence regulators in mycobacteria and plasmodium and, more recently, proposed a link between the tRNA epitranscriptome and translation adaptation during the CHIKV, DENV and SARS-CoV2 infections^26,27,29,50–53^. However, nothing was known concerning the relevance of the tRNA epitranscriptome in the context of IAV infection, and whether it also plays a role in regulating host antiviral responses. In this study, we uncovered a multilayered interplay between host tRNA biology, cellular stress responses, and IAV replication. By combining tRNA modification profiling, tRIBO-Seq, transcriptomics and functional perturbation of specific tRNA-modifying enzymes, we demonstrate that IAV rewires host translational and stress pathways in a time-dependent manner, selectively engages specific tRNA species on ribosomes, benefits from the host’s U34-modified tRNAs that decode IAV’s A/U-rich codon bias, and that the host tRNA epitranscriptome constitutes both a viral target and a determinant of antiviral defense (**Figure 9**).

**Figure 9.**
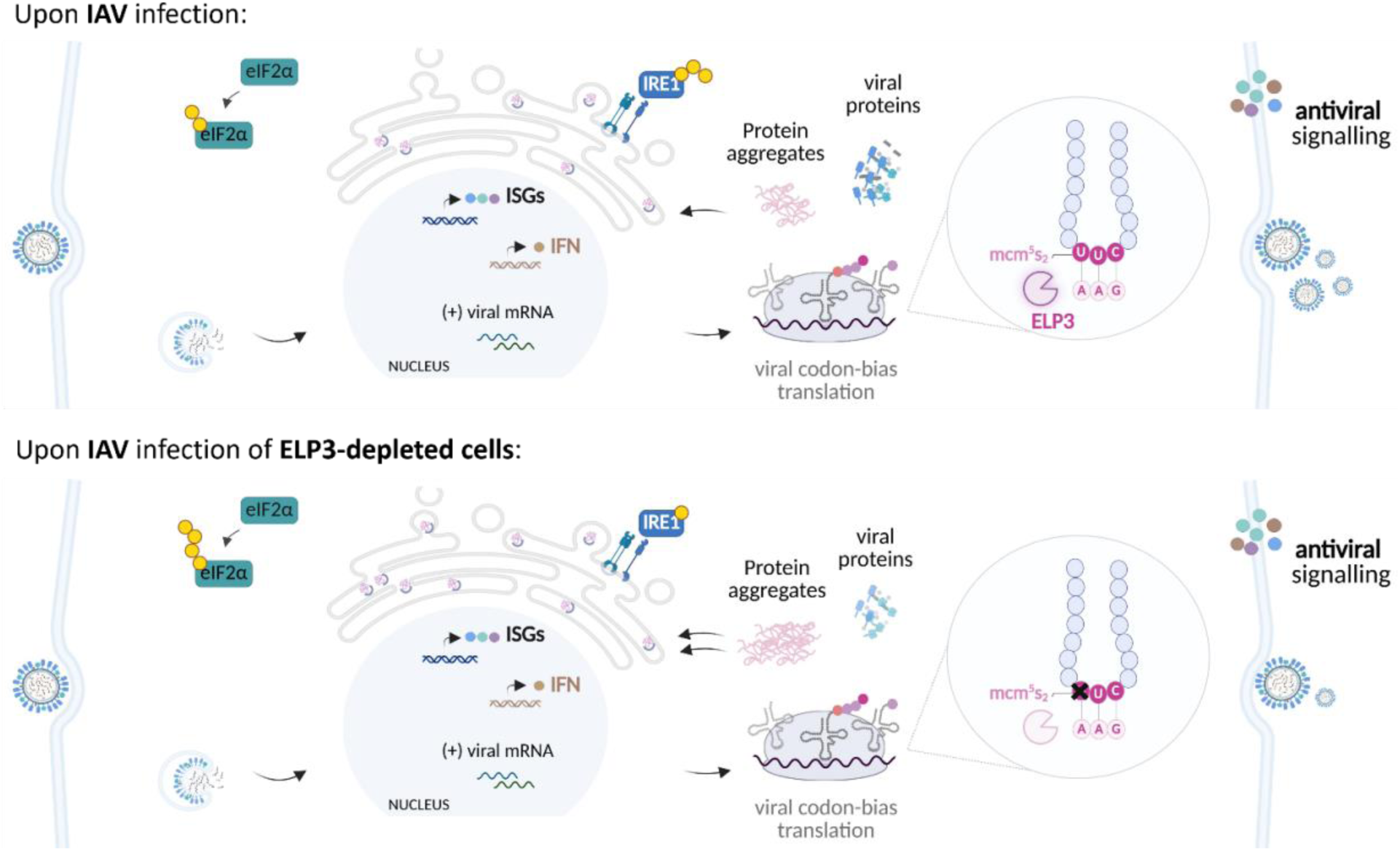
Proposed model -Wobble uridine modifications are reprogrammed at later stages of IAV infection. Under normal circumstances, the host cell tRNA modification machinery is activated to reprogram codon optimality during IAV infection. Wobble uridine modifications and its writers, including ELP3, are particularly important in this context as they boost the recognition of the sub-optimal IAV codons. Proteostasis disruption during IAV infection increases protein aggregation, triggering ER stress and the ISR-eIF2α and the UPR-IRE1α-XBP1 mechanisms. On the other hand, in environments lacking ELP3, tRNA hypomodifications are induced, impairing viral codon-biased translation, and intensifying proteostasis disruption. Specifically, the knockdown of ELP3 prior to IAV infection increases the phosphorylation of eIF2α and blocks the UPR-IRE1α-XBP1 arm, essential to IAV propagation. Created using Biorender.com

By exploring the dynamics of tRNA modifications within a single cycle of IAV infection, ranging from 2 to 12 hpi, we found evidence of time-dependent tRNA epitranscriptome modulation in response to infection. Several tRNA modifications, including ac⁴C, i⁶A, t⁶A, Q, m²,²G, and mcm⁵U, decline significantly at late infection stages. On the other hand, the downstream wobble modification mcm⁵s²U presents an opposite trend, accumulating at late infection timepoints, coinciding with enhanced thiolation of tRNA^Lys^_UUU_ and tRNA^Glu^_UUC_. These tRNAs decode AAA and GAA codons, two of the most enriched codons in the IAV genome, implying that IAV benefits from the host’s remodeling of U34 tRNA modification. In support of our observations, the reprogramming of mcm^5^U_34_ and mcm^5^s^2^U_34_ was also evident during CHIKV, Sars-CoV2 and DENV infection, particularly at later stages of infection. Specifically, during CHIKV infection, the levels of mcm^5^U_34_ increased upon viral infection, whereas the levels of mcm^5^s^2^U_34_ stayed constant^27^. On the other hand, and similar to our data, the levels of mcm^5^s^2^U_34_ during DENV infection were always increased when compared to non-infected controls and to the levels of mcm^5^U_34_ alone^26^. After SARS-CoV2 infection, fluctuations on mcm^5^U and mcm^5^s^2^U modification levels were found at different infection timepoints. For example, both modifications were increased at 6 hpi and at 24 hpi, whereas at 16 hpi and at 48 hpi mcm^5^U and mcm^5^s^2^U showed opposite trends, with increased mcm^5^s^2^U levels and decreased mcm^5^U levels when compared with non-infected controls^26,53^, similarly to our observations in IAV infected cells at 12hpi.

Transcriptomic analysis revealed that host gene expression shifts considerably at 8 and 12 hpi, coinciding with the peak of viral protein synthesis. In line with increased levels of mcm^5^s^2^U_34_, up-regulated genes at these late timepoints displayed a pronounced enrichment for A-ending codons, especially AAA, AGA, GAA, and CAA, which are decoded by mcm⁵s²U-modified U34 tRNAs and are overrepresented in the IAV genome. The convergence of codon usage remodeling and tRNA modification dynamics suggests a coordinated response in which both host and virus rely on a shared pool of U34-modified tRNAs during periods of high translational load. Consistent with this, tRIBO-seq revealed preferential loading of specific tRNA onto ribosomes, including tRNA^Lys^_UUU_ and tRNA^Glu^_UUC_, at late infection timepoints. Moreover, the tRNA ribosomal enrichment positively correlated with viral codon usage. These shifts occurred despite stable global tRNA levels, highlighting selective ribosome loading rather than changes in tRNA abundance, in line with what was observed previously for IAV and vaccinia virus^25^. Together, these results reveal that IAV infection contributes to rewire host translation in a codon-dependent manner, in which U34-modified tRNAs are preferentially mobilized, supporting viral translation.

The relevance of U34 tRNA modifications to viral decoding was further validated through loss-of-function studies. Silencing of ELP3, the catalytic subunit of the Elongator complex that initiates mcm⁵/mcm⁵s²U synthesis, led to significant hypomodification of mcm⁵U and mcm⁵s²U across the tRNA pool and caused a broad decline in IAV protein synthesis, viral genome decoding, and infectious particle production. This was expected as several reports have shown that tRNA hypomodifications, concomitantly to the knockdown of ELP3, compromise translation of A/U-rich transcripts, leading to increased protein aggregation, amino acid misincorporation and ribosome stalling, as well as activation of the ISR^15,47,54,55^. Importantly, transfection of a plasmid encoding tRNA^Lys^_UUU_ restored viral protein levels in ELP3-depleted cells, demonstrating that the antiviral phenotype derives directly from impaired U34 tRNA modification rather than from defects in tRNA abundance. These data pinpoint ELP3-dependent modifications as an essential host determinant of IAV translation efficiency, under our experimental conditions. Future assessment of codon-resolved translation under ELP3 deficiency will be important to further substantiate codon-specific decoding defects.

Silencing ELP3 did not alter the RIG-I/MAVS signaling, nor did it potentiate IFN or ISG expression during infection, indicating that the antiviral phenotype observed in ELP3-depleted cells is not attributable to increased innate immune sensing. On the other hand, ELP3 overexpression increased STAT1 phosphorylation in both infected and uninfected cells, suggesting a certain priming of the antiviral state independently of classical RIG-I/MAVS activation. This asymmetry reflected by absence of immune enhancement upon ELP3 loss but robust STAT1 activation upon ELP3 gain, highlights a non-canonical role for ELP3 in shaping cytokine signaling, possibly through translational or proteostasis-linked mechanisms. This is in line with previous studies showing that, in macrophages, deletion of ELP3 impairs the pattern recognition receptor (PPR)-IFN I-ISG axis, and that ELP3 is required for innate immune gene induction following infection with IAV^56^. Still, we did not define the ELP3 dose-response curve for viral replication and stress/immune outputs. Therefore, we cannot yet specify the optimal range of ELP3-dependent U34 modifications that maximizes IAV translation while minimizing antiviral priming. Additionally, we found that ELP3 mRNA and protein levels decline during mid-to-late infection correlates with increased ER stress and activation of the ISR potentiated by IAV infection. ELP3 knockdown prior to IAV infection hyperactivated the phosphorylation of eIF2α, an ISR marker, but hindered the IRE1-XBP1 branch of the UPR that is activated upon IAV infection^46,49^. Studies involving alphaarterivirus and ZIKV also show that, upon infection, the IRE1/XBP1s axis is preferentially activated^57,58^. Because IAV relies on the IRE1α–XBP1 pathway for productive replication, the inability to activate this UPR arm in the absence of ELP3 likely contributes to impaired viral growth. It is then reasonable to speculate that under tRNA hypomodification conditions, IAV-infected cells are trying to integrate diverse ER stress signals to modulate their activity. We hypothesize that the IRE1 pathway is less activated in detriment of eIF2α phosphorylation in environments of elevated ER stress that occur in the absence of ELP3. In support of this hypothesis, overexpression of tRNA^Lys^_UUU_ partially restored XBP1 splicing, further connecting tRNA modification status to the balance of host stress responses. Pharmacological induction of ER stress phenocopied the loss of ELP3. Similarly to what we had observed during IAV infection, ELP3 was differentially expressed following exposure of A549 cells to DTT and SA, two ER stress inducers. Of note, DTT exposure induced an ELP3 expression reduction of approximately 10-15%, similarly to what was observed upon IAV infection and that was sufficient to elicit XBP1 splicing. On the other hand, SA exposure led to a decrease of more than 50% of ELP3 expression and a suppression of XBP1 splicing, similarly to what occurs in IAV infected cells that were previously treated with an siELP3. This indicates that the reduced expression of ELP3 throughout IAV infection may hence reflect a broader cellular response to the proteotoxic stress induced by IAV, and that ELP3 expression is a titer for the UPR. It also suggests that viral ER stress feeds back onto the host tRNA modification machinery, creating a regulatory loop in which IAV-induced stress reduces ELP3 abundance and alters the translation landscape.

Importantly, we show that this overall strategy is not unique to IAV. VSV, another RNA virus with strong codon bias toward A/U-rich codons, also depends on ELP3 for efficient decoding and propagation. ELP3 depletion in VSV-infected cells reduced mcm⁵s²U levels, decreased viral protein production, impaired XBP1 activation, and reduced viral titers. These findings, together with previous studies in other RNA viruses namely DENV, CHIKV and SARS-CoV2, indicate that reliance on U34-modified tRNAs is a shared feature among RNA viruses with biased codon usage.

Altogether, our findings identify the tRNA epitranscriptome as a critical regulator of host translation and stress signaling during IAV infection. By reshaping codon usage patterns, modulating tRNA modification pathways, and perturbing the balance of ISR and UPR signaling, IAV exploits host translation control mechanisms to optimize viral protein synthesis. These results suggest that the tRNA epitranscriptome may represent a promising target for host-based antiviral therapy and suggest that codon-biased RNA viruses take advantage of the host tRNA modification networks to optimize its genome translation. Future studies dissecting the interplay between specific tRNA modifications, stress granule dynamics, and viral codon evolution may reveal additional strategies by which RNA viruses co-opt the host epitranscriptome for efficient replication. It will be important to evaluate the relevance of the other tRNA modifications that were also found altered during the IAV infection cycle, elucidate which specific tRNA isoacceptors are affected, and include proteome-wide and codon-resolved translation analyses. All in all, our combined data provide strong evidence on how RNA viruses exploit the host tRNA epitranscriptome landscape to not only fine-tune viral translation but also to modulate cellular stress and immune responses, highlighting the tRNA epitranscriptome as a multifunctional hub in virus–host interactions.

## Material and Methods

### Cell lines

The human adenocarcinoma alveolar basal epithelial (A549), Madin-Darby Canine Kidney (MDCK), human Embryonic Kidney 293T (HEK293T), and the monkey epithelial kidney (VERO) cells lines were cultured in Dulbecco’s Modified Eagle Medium high glucose (4,5 g/L) (DMEM) supplemented with 10% of fetal bovine serum (FBS, Sigma) and 1% penicillin/streptomycin solution (P/S) (all from GIBCO, Thermo Scientific, Waltham, MA, USA) at 37°C in a humidified 5% CO_2_ incubator. Cells were detached using TrypLE Express (Gibco) and resuspended in fresh media to count using a Neubauer chamber. All cell lines were frequently tested for mycoplasma contamination.

### Virus stocks preparation and plaque assay

#### Influenza A virus

Reverse-genetics derived influenza A/PR8/34 (IAV PR8; H1N1) was generated in HEK293T cells using a pDUAL plasmid system. HEK293T cells were transfected with eight plasmids, each encoding a gene segment of IAV PR8 (PB2, PB1, PA, HA, NP, NA, M, and NS1), using polyethylenimine (PEI) transfection reagent in DMEM high glucose (4.5 g/L) with pyruvate (0.11 g/L), supplemented with 1% glutamine (infection media). The following day, the medium was replaced by 2 mL of serum-free media (SFM) supplemented with 1% P/S, 0.14% bovine serum albumin (BSA) and 1 μg/mL of N-tpsyl-L-phenylalanine chloromethyl ketone (TPCK)-treated trypsin (1:1000). Forty-eight hours later, the supernatant containing a low virus count (P0) was collected, centrifuged at 3000 rpm for 5 minutes, and stored at -80°C for subsequent production of workable titter (P1) viruses. MDCK monolayers were infected with P0 aliquots in 5 mL of SFM supplemented with 0.14% bovine serum albumin (BSA) and 1 μg/mL Trypsin-TPCK, at a multiplicity of infection (MOI) of 0.01. After one hour, cells were overlaid with 10 mL of the above-mentioned media and cultured for an additional two days at 37 °C, 5% CO_2_. Viral supernatants were then collected after centrifugation, aliquoted, stored at -80°C, and later tittered using the plaque assay.

Plaque assays were conducted to determine viral titers as plaque forming units per milliliter (pfu/ml). MDCK monolayers were rinsed with phosphate buffered saline (PBS) and infected with 10-fold dilutions of viral supernatants diluted in infection media for 15 minutes on the rocker and 45 minutes at 37°C in 5% CO_2_. Subsequently, cells were overlaid with infection media supplemented with 0.14% of BSA, Trypsin-TPCK (1:1000) and 2.4% of Avicel and incubated at 37°C in 5% CO_2_ for approximately 1.5 days. Plaques on MDCK monolayers were washed with PBS and stained with 0.1% toluidine blue in 4% paraformaldehyde (PFA) solution on the rocker overnight. The next day, the wells were washed, air-dried and plaques were counted.

### Vesicular stomatitis virus

VSV Indiana stain parental stock was kindly provided by Dr. Jonathan Kagan (Boston Children’s Hospital, Harvard Medical School, Boston, MA, USA). Virus stocks were propagated in Vero monolayers, using a MOI of 0.001. The virus inoculum was diluted in SFM and after a one-hour virus adsorption period, an incubation of 18 h ensued. The clarified supernatant containing the viruses was centrifugated for 5 minutes at 4°C. Subsequently, a 90-minute ultracentrifugation (24000 rpm) was performed at 4°C. The supernatant was discarded, the pellet was diluted in 350 μL of NTE buffer (0.1M NaCl, 1mM EDTA, 0.1M Tris pH 7.4) and left to incubate on ice overnight. The following day, 4 mL of ice-cold sucrose 10% NTE was mixed with 1 mL of VSV suspension and subjected to a 60-minute ultracentrifugation (40500 rpm) at 4°C. The resulting pellet was incubated with 500 μL of NTE buffer on ice overnight. Finally, the virus suspension was aliquoted and stored at -80°C. To determine VSV titers, plaque assays were performed in Vero cells as previously described.

### Infection experiments

A549 cells were grown to 80% confluency and infected with IAV PR8 or VSV at a MOI of 3 in infection media and SFM, respectively, for 15 minutes with agitation at room temperature (RT) and for 45 minutes at 37°C and 5% CO2. Cells were overlaid with 20% FBS and incubated for the desired time-points, ranging from 2 to 12hpi, at 37°C and 5% CO_2_. For plaque assays, cells were infected as described. After one hour of virus absorption, the media was removed and cells were rinsed with acid buffer 1x (135 mM NaCl, 10 mM KCl, 40 mM citric acid, pH 3) and PBS. Afterwards, cells were incubated with infection media or SFM supplemented with 0.14% BSA, and 1 μg/mL Trypsin-TPCK (for IAV PR8 only), for up to 16hpi.

### Reverse transfection with siRNAs

A549 cells were reversely transfected with a siGenome SMARTpool human negative control siRNA (siCTR) and a siRNA targeting the tRNA-modifying enzyme ELP3 (siELP3) (Dharmacon) in 6-well plates. Briefly, master mixes for each transfection condition were prepared containing 48 μL of a 500 nM siRNA duplex, 352 μL of Opti-MEM Reduced Serum Media (Alfagene) and 4 μL/well of lipofectamine mix (Dharmacon). After a 30 minutes incubation, 1.5x10^5^ cells/mL were added to each well in culture media without antibiotics and incubated for 48 hours at 37°C in a CO_2_ incubator.

### ELP3 overexpression

A549 cells were transfected with the ELP3-expressing plasmid (pELP3) (OriGene, ELP3 (NM_018091) Human Untagged Clone) or with an empty control plasmid (pMOCK) (OriGene, pCMV6-AC Mammalian Expression Vector). Transfections were performed using TurboFect™ Transfection Reagent (Thermo Fisher Scientific, USA) in antibiotic-free medium, following the manufacturer’s instructions, and carried out in 12-well plates. Prior transfection, cells were detached from a culture plate using TrypLE Express Enzyme (Thermo Fisher Scientific, USA). The cell suspension was centrifuged at 3000 rpm for 4 minutes, and the supernatant was discarded. The resulting pellet was resuspended in 1 mL of antibiotic-free medium. Cell density was determined using a Neubauer counting chamber. Based on the cell count, a final suspension was prepared at a concentration of 12×10^4^ cells/mL in antibiotic-free medium and homogenized by gentle inversion. Subsequently, 1 mL of the cell suspension was plated into each well of a 12-well plate. The plates were then incubated for 24 hours at 37°C in an atmosphere with 5% CO_2_. After 24 hours of incubation, a transfection mixture was prepared by combining the appropriate plasmid (0.5 ug/mL; pELP3 or pMOCK), TurboFect™ Transfection Reagent (Thermo Fisher Scientific, USA), and Gibco Opti-MEM Reduced Serum Medium (Thermo Fisher Scientific, USA). This mixture was incubated at room temperature for 15 minutes. In the 12-well plate, approximately 700 μL of the medium was removed from each well. Subsequently, 200 μL of the transfection mixture was added dropwise to each well. The plates were returned to the 37°C incubator. After 24h of incubation, the medium was removed and IAV infection was performed as described above. Infected and mock-infected cells were collected at the desired infection time-points for further analysis.

### SunSET assay

SunSET assay was performed as previously reported ^41^. Briefly, to determine protein synthesis rate, upon reverse transfections with siRNAs, cells were incubated with puromycin (1 μg/ml) for 15 minutes at 37°C in 5% CO_2_. Cells were detached, centrifugated and washed with PBS before proceeding to protein extraction for Western Blot analysis. An anti-puromycin antibody was used (clone 12D10; Sigma-Aldrich, MABE343, 1:10000 dilution).

### RNA extraction

Total RNA used in LC/MS-MS experiments and qPCR was extracted using TRIsure™ (Bioline) according to the manufactureŕs instructions. For the tRNA and RNA sequencing experiments, enriched small and large RNA fractions were isolated using the mirVana^TM^ miRNA isolation kit (Ambion, Waltham) in combination with the Zymo RNA Clean and concentrator kit (Zymo Research). Briefly, following cell lysis, initial RNA phenolic extraction was performed using the miRVana^TM^ kit, following the miRVana^TM^ Organic Extraction step according to the manufacturer’s protocol. The obtained aqueous phase was then further processed to obtain enriched small RNA and large RNA fractions using the Zymo RNA Clean and concentrator kit and following the manufacturer’s protocol.

### cDNA synthesis and qRT-PCR

Five hundred nanograms of RNA were used for cDNA synthesis using the Applied Biosystems^TM^ High-Capacity RNA-to-cDNA^TM^ Kit (Thermo Fisher Scientific), following the manufacturer’s instructions. The obtained cDNA was used for qPCRs using 2x SYBR Green qPCR Master Mix Low Rox (Bimake, Houston, TX, United States) and the reactions were carried out in the Applied Biosystems 7500 Real-Time PCR System (Applied Biosystems, Wathman, Massachusetts, USA). cDNA samples were diluted 1:10 and the concentration of each primer was 250 nM, with the exception for IFN-β in which we use undiluted samples and a primer concentration of 500 nM. The primer sequences used for XBP1 were 5- TTACGAGAGAAAACTCATGGCC’-3’ and 5’- GGGTCCAAGTTGTCCAGAATGC-3’; for sXBP1 were 5’- CTGAGTCCGAATCAGGTGCAG-3’ and 5’-ATCCATGGGGAGATGTTCTGG-3’; for IFN-β were 5’-TGGCACAACAGGTAGTAGGC-3’ and 5’-TGGAGAAGCACAACAGGAGAG-3’; for IRF1 were 5’- ATGCCCTCCACCTCTGAAG-3’ and 5’-CCACTCCGACTGCTCCAA-3’; for IFN-λ were 5’- GAA GCC TCA GGT CCC AAT – 3’ and 5’ - GCA GGT TCA AAT CTC TGT CAC – 3’; for GAPDH (used as internal control) were 5’-AAGGTGAAGGTCGGAGTC-3’ and 5’-GGGTGGAATCATATTGGAACAT-3’ (Eurofins Genomics, Ebersberg, Germany). Expression of ELP3, URM1, and GAPDH was evaluated using Taqman probes and the TaqMan^TM^ Gene Expression Master Mix (Thermo Fisher Scientific) following the manufacturer’s guidelines. Data analysis was performed using the quantitative 2−ΔΔCT method.

### Isolation of tRNAs by HPLC

tRNAs were separated from total RNA samples by size exclusion chromatography (SEC), using an Agilent 1100 HPLC system equipped with an AdvanceBio column 300 Å column (Agilent, Waldbronn). tRNA peaks were uncovered by measuring absorbance at 260 nm and collected based on pre-established retention times. tRNAs were concentrated by SpeedVac (Thermo Fisher Scientific) and precipitated in 100% ethanol and 0.5M ammonium acetate overnight at -20°C. Following a 30-minute centrifugation at 4°C, the pelleted tRNAs were rehydrated with milliQ water and quantified in a nanophotometer (NP80, IMPLEN).

### Quantification of tRNA modifications by LC-MS/MS

Two hundred nanograms of the purified tRNA samples were digested into nucleosides for 2h at 37°C with benzonase (2U/μL), phosphodiesterase I (0.2 U/μL), alkaline phosphatase (2 U/μL), Tris pH 8.0 (5 mM), magnesium chloride (1 mM), tetrahydrouridine (5 μg), butylated hydroxytoluene (10 µM) and pentostatin (1 µg). For quantitative mass spectrometry of the nucleosides, an Agilent 1290 Infinity II equipped with a diode-array detector (DAD) combined with an Agilent Technologies G6470A Triple Quad system and electrospray ionization (ESI-MS, Agilent Jetstream) was used. Operating parameters: positive-ion mode, skimmer voltage of 15 V, cell accelerator voltage of 5 V, N2 gas temperature of 230 °C and N2 gas flow of 6 L/min, N2 sheath gas temperature of 400 °C with a flow of 12 L/min, capillary voltage of 2500 V, nozzle voltage of 0 V, nebulizer at 40 psi. The instrument was operated in dynamic multiple reaction monitoring mode (dMRM).

For separation a Synergi, 2.5 µm Fusion-RP, 100 Å, 100 x 2 mm column (Phenomenex, Torrance, California, USA) at 35 °C and a flow rate of 0.35 mL/min was used. Mobile phase A consisted of 5 mM aqueous NH4OAc buffer, brought to a pH of 5.3 with glacial acetic acid (65 µL/L) and mobile phase B consisted of organic solvent acetonitrile (Roth, Ultra-LC-MS grade). The gradient started at 100 % A for 1 min and an increase of 10% B over a period of 4 min afterwards. B was then increased to 40 % for 2 min and maintained for 1 min before returning to 100% A over a period of 0.5 min followed by a re-equilibration period for 2.5 min.

For external calibration solutions were prepared using synthetic nucleosides. The calibration solutions ranged from 0.025 to 100 pmol for canonical nucleosides and from 0.00125 pmol to 5 pmol for modified nucleosides, except of pseudouridine which ranged from 0.005 to 20 pmol. 10 µL of each sample and calibration was injected into the LC-MS system for analysis. Additionally, 1 µL of previously prepared and digested stable isotope labeled internal standard was co-injected.

Raw data was analyzed using quantitative MassHunter Software from Agilent.

### RNA-Sequencing

In total, 18 RNA samples (3 biological replicates corresponding to the 2hpi, 8hpi, 12hpi and respective mock infected samples) were sent for transcriptomic analysis. All RNA sequencing procedures were conducted by Biomarker Technologies (BMKGENE, Germany).

A combination of miRVana and Zymo clean RNA up kits were used to ensure phenolic extraction of the lysates and enrichment in small non-coding RNA fraction and in long RNA fraction, as described above. The long RNA fraction was used to proceed with RNA-sequencing. Purity, concentration and integrity of the RNA samples were examined by Nanodrop (Thermo Fisher Scientific). RNA integrity was also assessed using the RNA Nano 6000 Assay Kit of the Agilent Bioanalyzer 2100 system (Agilent Technologies) Only RNA with RINs above 7 were sequenced, resulting in a total of 17 samples sequenced, as one of the 2hpi mock did not pass the quality threshold.

For library preparation, 1μg RNA per sample was used as input material. Sequencing libraries were generated using NEBNext UltraTM RNA Library Prep Kit for Illumina. mRNA was isolated by Oligo(dT)-attached magnetic beads and then randomly fragmented in fragmentation buffer. First-strand cDNA was synthesized with fragmented mRNA as template and random hexamers as primers, followed by second-strand synthesis with addition of PCR buffer, dNTPs, RNase H and DNA polymerase I. Purification of cDNA was processed with AMPure XP beads. Double-strand cDNA was subjected to end repair. Adenosine was added to the end and ligated to adapters. AMPure XP beads were applied here to select fragments within a size range of 300-400 bp. cDNA library was obtained by PCR on the cDNA fragments generated. PCR products were purified (AMPure XP system) and library quality was assessed on the Agilent Bioanalyzer 2100 system. The qualified library was clustered on a cBot Cluster Generation System using TruSeq PE Cluster Kit v4-cBot-HS (Illumina) according to the manufacturer’s instructions. After, the library preparations were sequenced on an Illumina platform and paired-end reads were generated.

The 17 samples sequenced generated 113.68 Gb Clean Data. At least 6.02 Gb clean data were generated for each sample with minimum 96.89% of clean data achieved a quality score of Q30. Clean reads of each sample were mapped to the human reference genome. Mapping ratio ranged from 64.86% to 98.46%. All bioinformatic analyses were performed on the BMKCloud platform. Processing of raw data was done through in-house perl scripts. Briefly, in this step, raw data passed through several quality steps (removing adapter contaminations and reads with low quality) to obtain clean and high-quality data. Quality scores (which represents the possibility (P) of an incorrect base incorporation) were calculated. Nucleotide distribution on reads and GC content were also analyzed. Differentially expressed genes (DEGs) were identified using the criteria of Fold Change≥1.5 and Pvalue<0.05. Then GO/KEGG enrichment, GSEA, and protein interactions of DEGs were analyzed.

### Nano-tRNA-seq

Quantification of tRNA abundances was performed by nanopore sequencing as previously described^37^. The fraction containing ≤ 200 nt RNAs was obtained using the mirVana^TM^ miRNA isolation kit (Ambion, Waltham) following the manufacturer’s guidelines. In brief, samples were disrupted in a lysis/binding solution and treated with a miRNA homogenate additive for 10 minutes on ice. Afterwards, an Acid-Phenol:Chloroform solution was added and mixed before centrifugation at maximum speed for 10 minutes. The RNA-containing aqueous phase was removed to a new tube and precipitated with an 100% ethanol solution. The precipitated RNA was applied to a filter cartridge, centrifugated, and the flow-through was discarded. After washing steps, the RNA was eluted in RNAse-free water and quantified in a DeNovix DS-11 spectrophotometer. RNA integrity was evaluated using a 4200 TapeStation System. The purified tRNAs from IAV-infected and non-infected A549 cells were deacylated in 95 μL of 100 mM Tris-HCl (pH=9) at 37°C for 30 minutes. The Deacylated tRNAs were recovered using the Zymo RNA Clean and Concentrator-kit (Zymo Research, R1016) to retain RNAs ≥17 nt, following the manufacturer’s instructions. tRNA libraries were prepared using the Nano-tRNAseq method^37^. Briefly, custom barcoded RTA adapters were used in place of the standard ONT RTA to enable sample multiplexing. After reverse transcription, samples were pooled and purified using 1.8× AMPure RNAClean XP beads prior to sequencing adapter ligation. Total tRNA libraries were prepared using the ONT SQK-RNA002 kit, and sequenced in R9.4.1 flowcells, following standard ONT protocols.

### tRIBO-Seq

Ribosomes were isolated using the RiboLace kit (Immagina Biotechnology S.r.l., cat. no: GF001-12), according to the manufacturer’s instructions as indicated in Monti *et al*^38^, with some minor modifications. after isolation of tRNAs attached to the ribosomes using the RiboLace Pro kit (Immagina). Briefly, ∼5 million cells were used for each condition. Translation was arrested by addition of 10 ng/uL cycloheximide for 10 min at 37 °C, preserving ribosome–RNA complexes. Cell lysates were subjected to nuclease digestion using 5X higher nuclease concentration than recommended by the manufacturer, as we observed that the amount required to have complete digestion to monosomes has to be adapted in a cell-type specific manner, Recovered RNA was then quantified at 260 nm using a Nanodrop ND-1000 UV-VIS Spectrophotometer, and RNA integrity and size were assessed via Agilent 4200 TapeStation RNA HS ScreenTape Assay (cat. no: 5067-5579). Ribo-embedded tRNAs were then sequenced using Nano-tRNAseq method^37^ and SQK-RNA004 chemistry (ONT), with minor variations. Ribo-tRNA libraries were prepared using SQK-RNA004 kit and sequenced on MinION devices using FLO-004RA (FLO-MIN106).

### Nano-tRNA-seq: read processing and alignment

*Basecalling and mapping:* Basecalling was performed using dorado (v7.2.13, https://github.com/nanoporetech/dorado) using the HAC model for total tRNAs and the FAST model for ribo-tRNAs. Demultiplexing was conducted using SeqTagger v1.0d (https://github.com/novoalab/SeqTagger,^59^). Reads were aligned using mappy (v2.28) with the following parameters: -ax splice -k7 -w3 -n1 -m13 -s30 -A2 -B1 -O2,32 -E1,0 --secondary=no. Alignments were performed against a reduced reference genome comprising curated tRNA sequences (available at https://github.com/mimonti/IAV_tRIBOseq). Reads with mapping quality (MapQ) <10 were discarded.

*Filtering:* For downstream analyses, reads lacking the 3′ splint adapter were discarded, as those most likely correspond to mismapped reads. Full-length reads were defined by their start positions relative to the reference tRNAs. Reads containing full or partial 5′ splint adapters were kept for downstream analyses, and were obtained by applying the ‘filtering’ module in AMaNITA (v0.2.5, https://github.com/novoalab/AMaNITA/tree/master).

*Differential tRNA abundance and Principal Component Analysis (PCA)*: Differential tRNA abundance and principal component analysis (PCA) were performed using DESeq2 (v1.42.1,^60^). Raw count data were normalized using DESeq2 size-factor normalization. Differential abundance analysis was conducted separately for each time point and sequencing modality (Ribo-embedded and Total) by subsetting the dataset per time point. Within each subset, comparisons between Control and Infected samples were modeled using a design formula including treatment group and sequencing batch (∼ Group + Batch) to account for batch effects during statistical testing. P-values were adjusted for multiple testing using the Benjamini–Hochberg procedure^61^. For PCA, variance-stabilized data were generated using the varianceStabilizingTransformation function from DESeq2. To further correct for batch effects in visualization, batch-corrected expression values were obtained using the removeBatchEffect function from the limma package, incorporating the treatment design. PCA was then performed on the batch-corrected, variance-stabilized data using the prcomp function, and visualized using ggplot2. Notebooks and outputs for differential expression, modification analysis, and codon correlation analyses are available at https://github.com/mimonti/IAV_tRIBOseq.

### Codon-anticodon correlation analysis

Codon usage bias values for influenza A virus (IAV) were obtained from published relative synonymous codon usage (RSCU) tables^22^. Codons were matched to their cognate tRNA anticodons according to standard codon-anticodon pairing rules to generate a codon-tRNA mapping. tRNA differential expression was quantified from ribosome-associated (ribo-tRNA) and total tRNA datasets using DESeq2, as described above (see *Nano-tRNA-seq: data analysis*). Codon usage bias was defined as the difference between H1N1pdm strain and human cells (HC) values. To assess the relationship between viral codon usage and host tRNA regulation, codon usage bias was integrated with tRNA differential expression data based on anticodon identity. Pearson correlation coefficients (r) were calculated between codon usage bias and tRNA log2 fold change for each condition using complete observations. Statistical significance was assessed using two-sided Pearson correlation tests.

### Data availability

RNA-Seq raw sequencing data is deposited in Gene Expression Omnibus (GEO) under the accession number GSE328283.

Nano-tRNA-seq and tRIBO-seq raw sequencing data (fast5/pod5) and aligned BAM files have been deposited in the European Nucleotide Archive (ENA) under accession number **PRJEB111033.** All datasets are provided in demultiplexed format. Barcode assignments for each accession are detailed in Supplementary Table 2.

### Total protein extraction and Western blotting

Cells were detached, washed with PBS, and resuspended in Empigen Lysis Buffer (ELB) (0.5% Triton X-100, 50 mM HEPES, 250 mM NaCl, 1 mM DTT, 1 mM NaF, 2 mM EDTA, 1 mM EGTA, 1 mM PMSF, 1 mM Na3VO4 supplemented with a cocktail of protease inhibitors (Complete, EDTA-free, Roche). After sonication, the lysates were centrifugated for 20 minutes at 16.000g, the supernatants were collected, and the protein content was quantified using the BCA Protein Assay Reagent (Thermo Fisher Scientific) according to the manufacturer’s instructions. Protein lysates were transferred to nitrocellulose membranes, blocked with 5% non-fat dry milk diluted in tris-buffered saline (TBS)-tween and immunoblotted with antibodies against ELP3 (ThermoFisher, 1:200 dilution), anti-puromycin antibody (clone 12D10; Sigma-Aldrich, MABE343, 1:10000 dilution), anti-NP (ab20343, Abcam, 1:1000 dilution), anti-PB1 (Santa Cruz Biotechnology, 1:1000 dilution), anti-PB2 (Santa Cruz Biotechnology, 1:1000 dilution), anti-HA (clone C102, Santa Cruz Biotechnology, 1:1000 dilution), anti-M1 (FluAc, Santa Cruz Biotechnology, 1:1000 dilution), anti-M2 (clone 14C2, Santa Cruz Biotechnology, 1:1000 dilution), anti-NS1 (Santa Cruz Biotechnology, 1:1000 dilution), anti-IRE1α (clone 14C10 (3294), Cell Signaling Technology, 1:1000 dilution), anti-eIF2α (clone D-3, Santa Cruz Biotechnology, 1:1000 dilution), anti-phospho-eIF2α (clone S1, Abcam, 1:1000 dilution), anti-ATF4 (Bioworld, 1:1000 dilution), anti-phospho-ATF4 (Bioworld, 1:1000 dilution), anti-GADD34 (Invitrogen; 1:1000 dilution), anti-GAPDH (Cell signalling, 1:1000 dilution), anti-actin (Cell signalling, 1:5000 dilution), anti-vinculin (ProteinTech, 1:5000 dilution), β-tubulin anti-mouse (Life Technologies, 1:5000 dilution), and β-tubulin anti-rabbit (ProteinTech, 1:5000 dilution) overnight at 4°C. The secondary antibodies IRDye 680® Goat anti-rabbit or IRDye® 800 CW Goat anti-mouse (LI-COR Biosciences, 1:10000 dilution) were used for signal detection in an Odyssey Infrared Imaging system (LI-COR Biosciences).

### tRNA^Lys^_UUU_ overexpression

500 ng of a DNA plasmid carrying the tRNA^Lys^_UUU_ gene, obtained as previously described in ^15^, was transfected into siCTR- and siELP3-treated cells using Turbofect (Thermo Fisher), following manufacturer’s instructions. As a control, cells were also transfected with an empty vector using the same conditions. 24 hours later, cells were infected with IAV PR8 and collected at the desired time-points.

### Northern Blotting

1 μg of total RNA was electrophoresed on 10% polyacrylamide/urea gel. For gels containing APM, APM was added at a concentration of 20 μM in the gel mixture and 5 μg of total RNA was electrophoresed, as described in ^36^. Briefly, the three-layer APM gel consisted of a bottom Gel Layer of a 10% polyacrylamide gel containing 8 M urea without APM, a Middle Gel Layer of a 10% polyacrylamide gel containing 8 M urea with 100 µg/mL of APM (obtained from an APM stock concentration of 1 mg/mL in formamide), and a top gel layer of a 10% polyacrylamide gel containing 8 M urea without APM. After electrophoresis, for both regular northern blots and APM-Northern blots, RNA was transferred to a positively charged nylon membrane (Amersham Biosciences, Cat. RPN203B) using a semi-dry blotting system. The RNA was crosslinked to the membrane in a Stratalinker equipment at 1200 mJ/cm^2^ for 1 minute on each side. Membranes were blocked in hybridization buffer (50x Denhardt’s solution, 10% SDS, 20x SSPE and H_2_O) for 2 hours. Afterwards, membranes were incubated with the respective non-radioactive, fluorescently labeled probe overnight with agitation. Of note, both incubations were performed at the same conditions, 5°C lower than the melting temperature of each probe. In the next day, the membranes were washed four times, and the signal was detected using an Odyssey Infrared Imaging System.

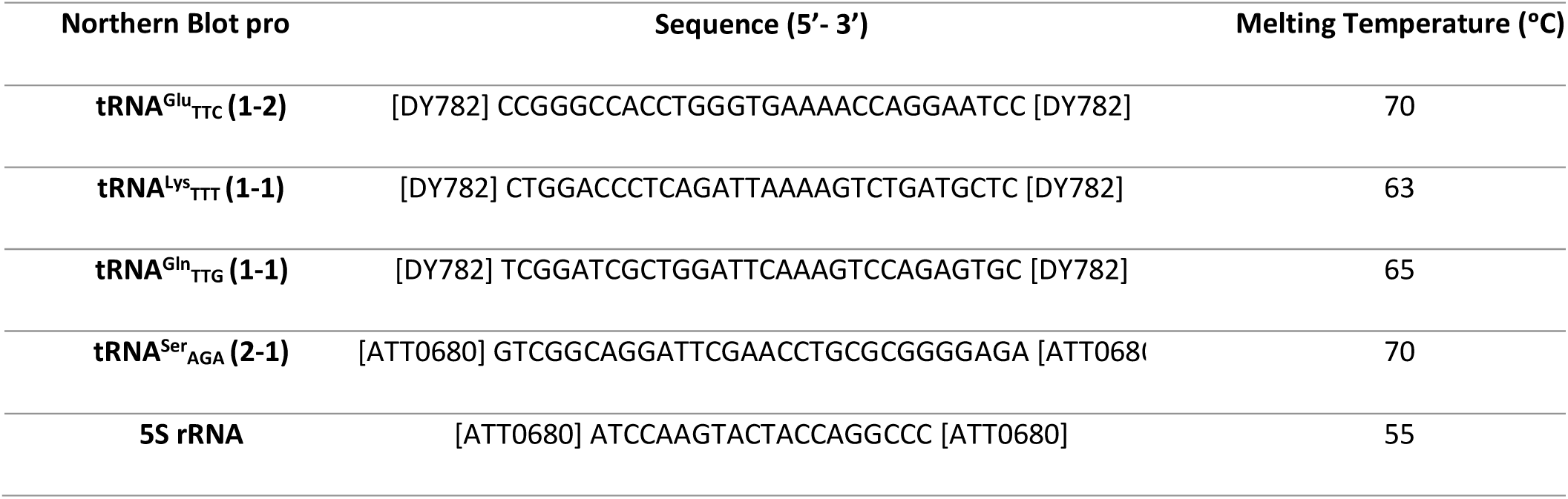

### Statistical analysis

Statistical analysis, except for sequencing experiments for which statistical analysis is explained in the corresponding methods section, was performed in the GraphPad Prim® v7 software, using Student’s unpaired t-test with Welsch correction or the one-way ANOVA test. The results were presented as mean values of the number of experiments. Error bars reflect standard deviation.

## Acknowledgements

We thank Catarina Almeida (for kindly providing anti-GADD34 antibodies, sodium arsenite, and XBP1 primers), Bruno Neves (for kindly providing anti-IRE1, and anti-phosphorylated IRE1α antibodies), and Hélio Albuquerque for synthetizing the APM component. We thank Xanthi-Lida Katopodi for granting beta access to the AMaNITA repository and for assistance with the analysis of tRIBO-seq and nano-tRNA-seq data. We are furthermore grateful to Leszek Piotr Pryszcz for providing scripts for basecalling, demultiplexing, and read mapping.

## Funding

This research was funded by the Portuguese Foundation for Science and Technology (FCT), POCH, FEDER, COMPETE2020, and COMPETE2030 through the grants SFRH/BD/146703/2019, COVID/BD/153557/2024, POCI-01-0145-FEDER-016630, POCI-01-0145-FEDER-029843, POCI-01-0145-FEDER-031378, COMPETE2030-FEDER- 00758600, UI/BD/151372/2021, UIDB/04501/2025, under the scope of the Operational Program “Competitiveness and internationalization”, in its FEDER/FNR component. It was also supported by the European Union through the Horizon 2020 program: H2020-WIDESPREAD-2020-5 ID-952373 and through the European Research Council under the grant agreement number 101042103 and 101187456 (to E.M.N). SK was funded by the Deutsche Forschungsgemeinschaft SFB1177 (259130777-SFB1177). M.M is supported by a Juan de la Cierva Postdoctoral fellowship (JDC2023-050553-I funded by MICIU/AEI /10.13039/501100011033 and FSE+).

## Author contributions

Conceptualization, D.R.R., D.R., and A.R.S.; methodology, D.R.R., A.N, M.M., L.L., M.P., M.B., K.K., E.M.N., S.K., D.R., and A.R.S.; data curation, A.R.G., D.R.R. and A.R.S.; writing-original draft preparation, D.R.R., M.M., A.R.S.; writing-review and editing, S.K., E.M.N., D.R., and A.R.S.; visualization, D.R.R., M.M., A.R.S.; supervision, D.R., A.R.S.; project administration, D.R., and A.R.S.; funding acquisition, A.R.S., D.R., E.M.N., and S.K. All authors have read and agreed to the published version of the manuscript.

## Disclosure statement

MM has received travel bursaries from Oxford Nanopore Technologies (ONT) to present her results at research conferences. EMN has received travel and accommodation expenses to speak at ONT conferences. EMN is listed as inventor in the patent describing the Nano-tRNAseq method (PCT/IB2023/059599, publication WO2024/069467). The remaining authors declare that they have no conflict of interest.

## Supplementary Material

**Supplementary table 1.**
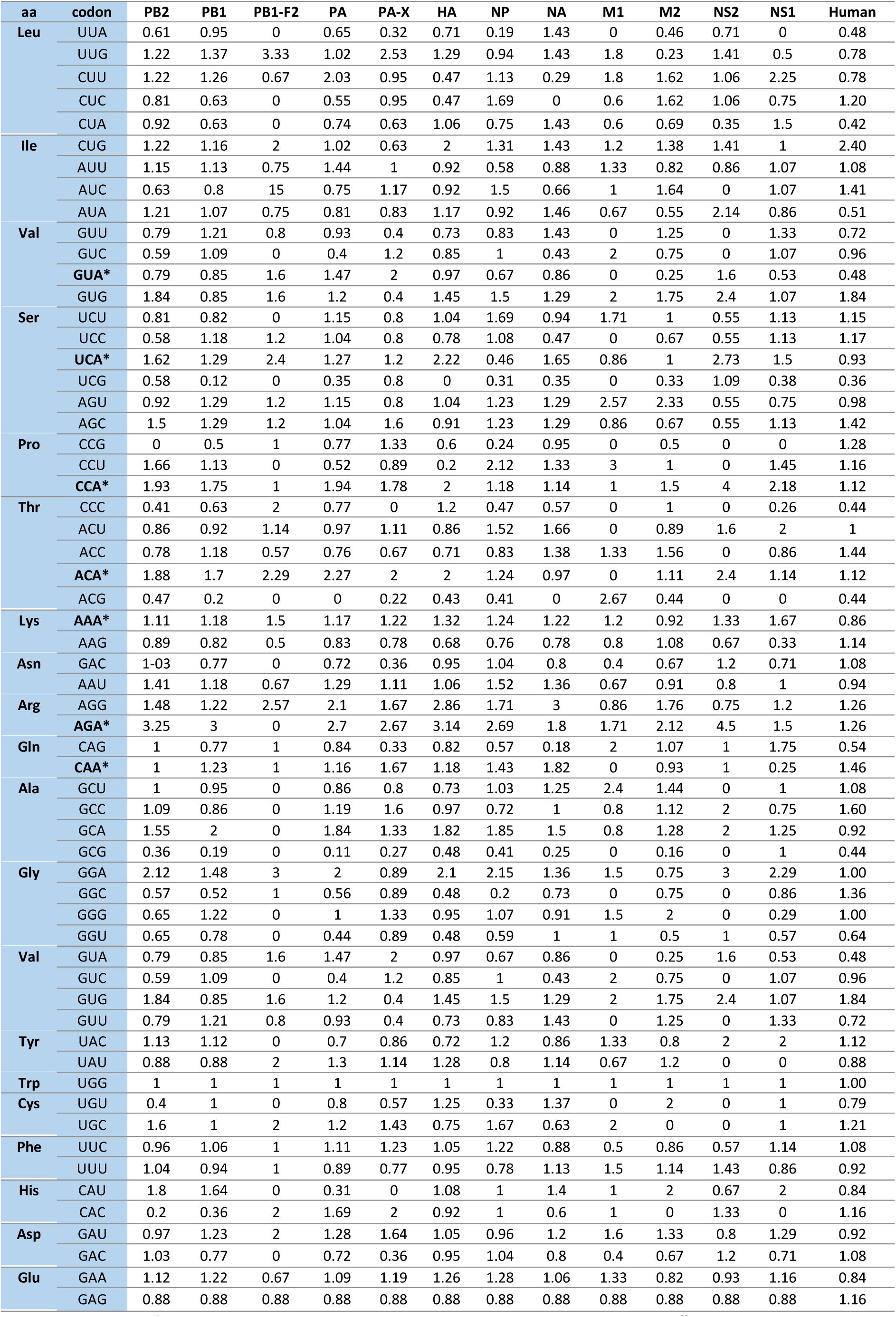
RSCU analysis of IAV complete codon sequences using the ANACONDA software. ^62^.

**Supplementary table 2.**
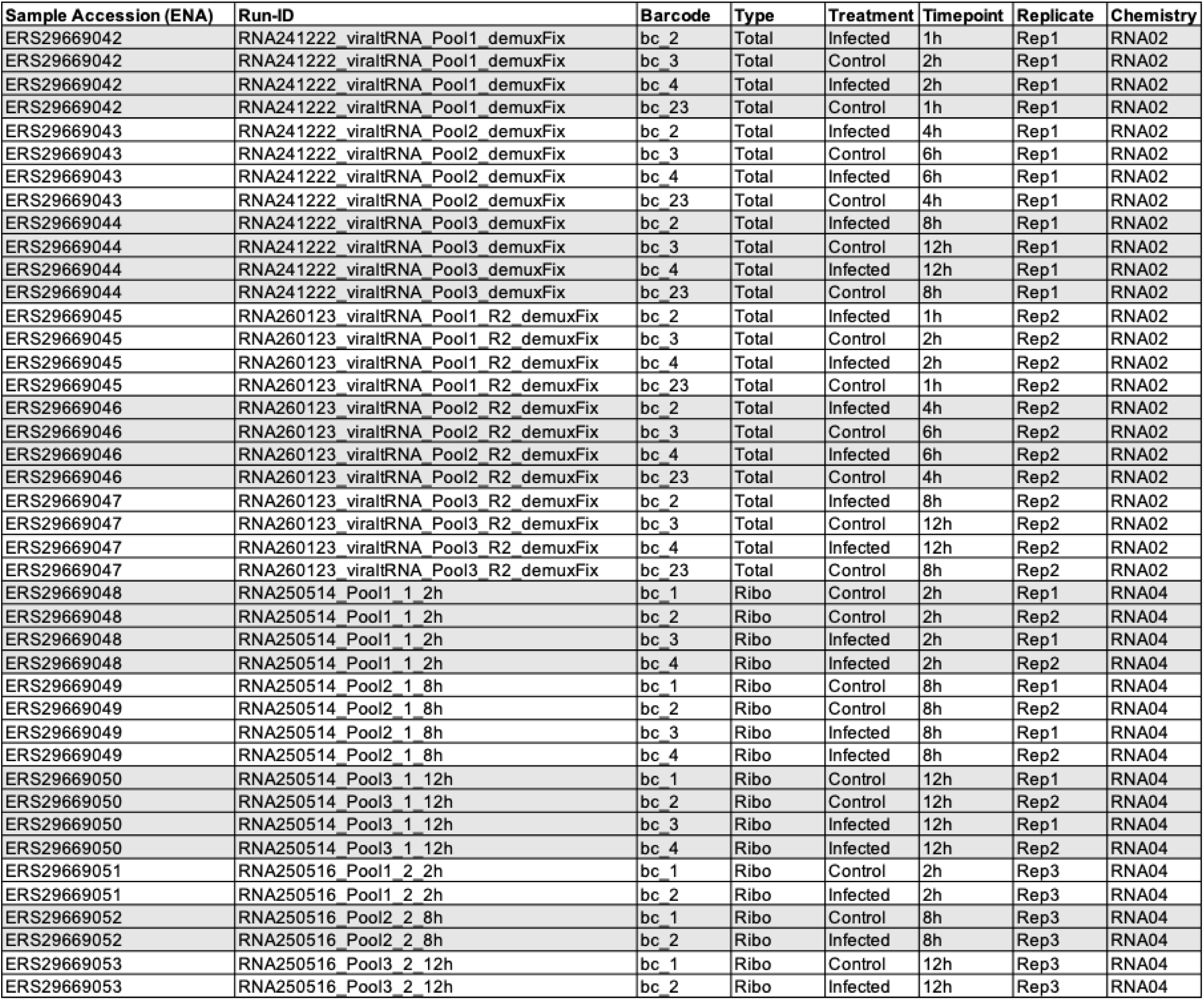
Barcode assignments for each ONT sample.

**Supplementary figure 1.**
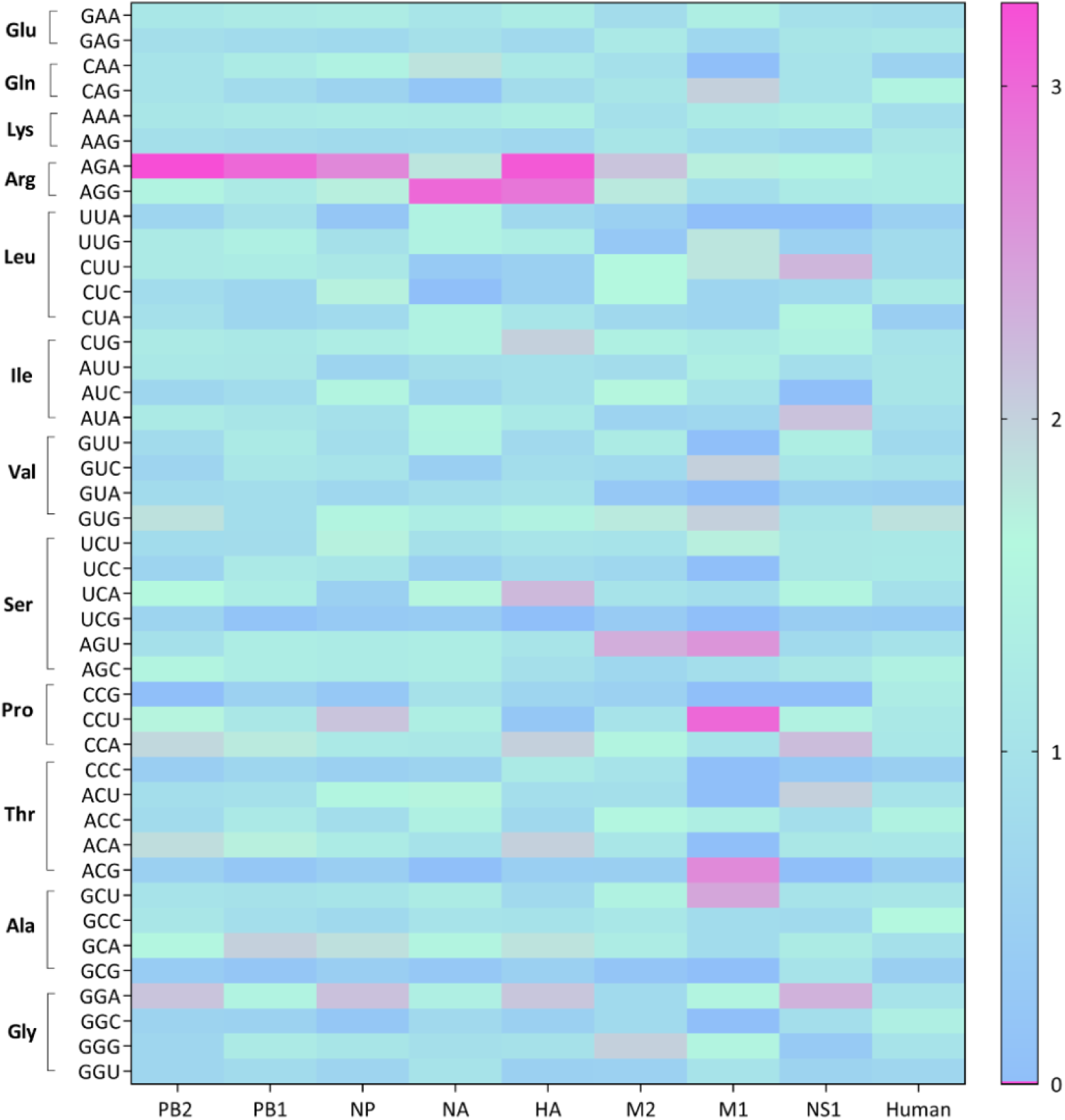
Heat map analysis showing codon usage frequencies of the complete coding sequences of IAV. Blue colors represent lower frequencies (under-represented codons) whereas pink colors represent higher frequencies (over-represented codons)

**Supplementary figure 2.**
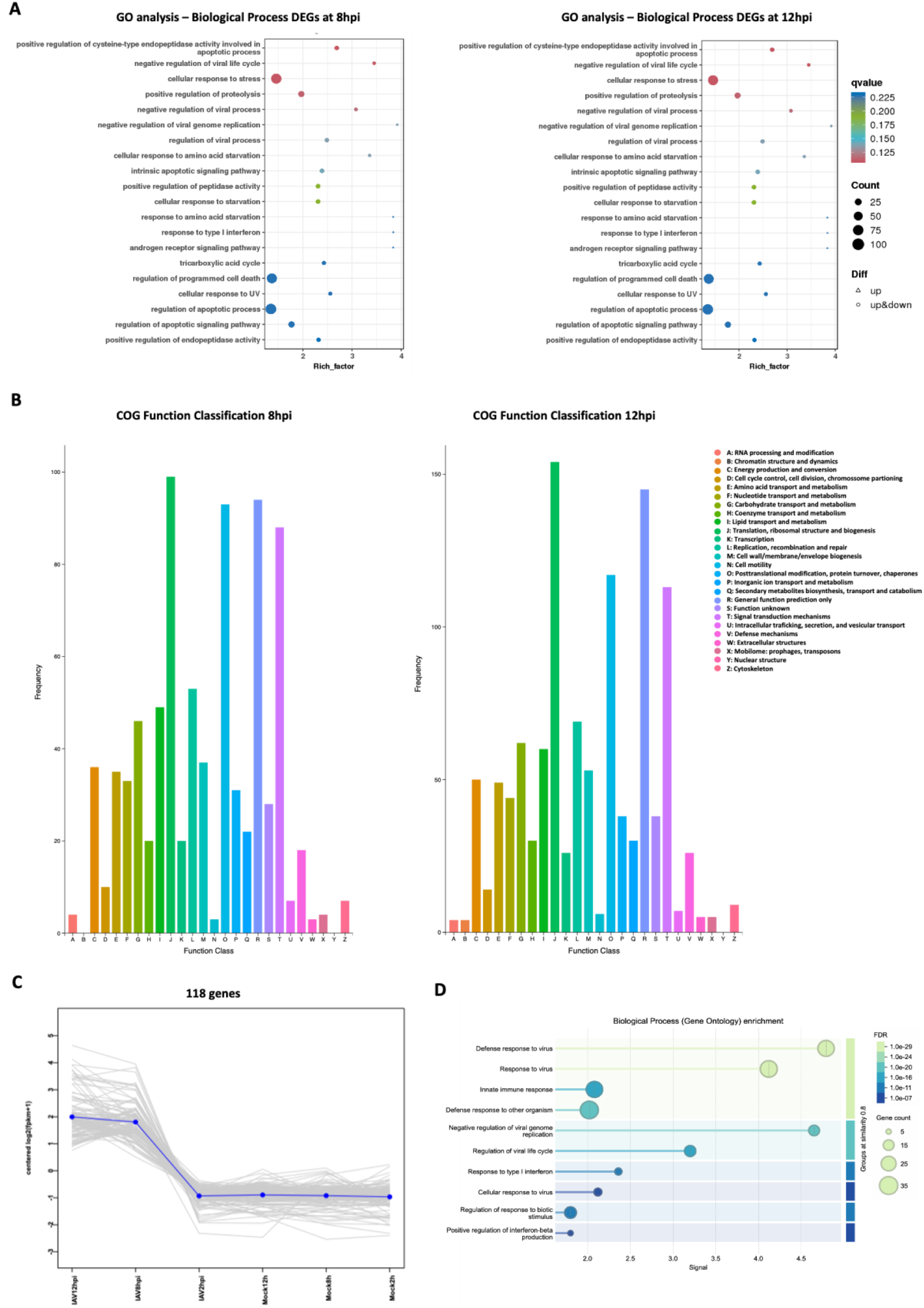
(A) GO analysis of the main biological processes affected at late infection timepoints 8 and 12 hpi. (B) COG analysis pf DEGs 8hpi and 12hpi. (C) WGCNA identifies a cluster of 118 genes that differentiate IAV 8 hpi and IAV 12 hpi samples from the remaining samples. (D) GO enrichment analysis (Biological function) of the 118 genes identified in the cluster.

**Supplementary Figure 3.**
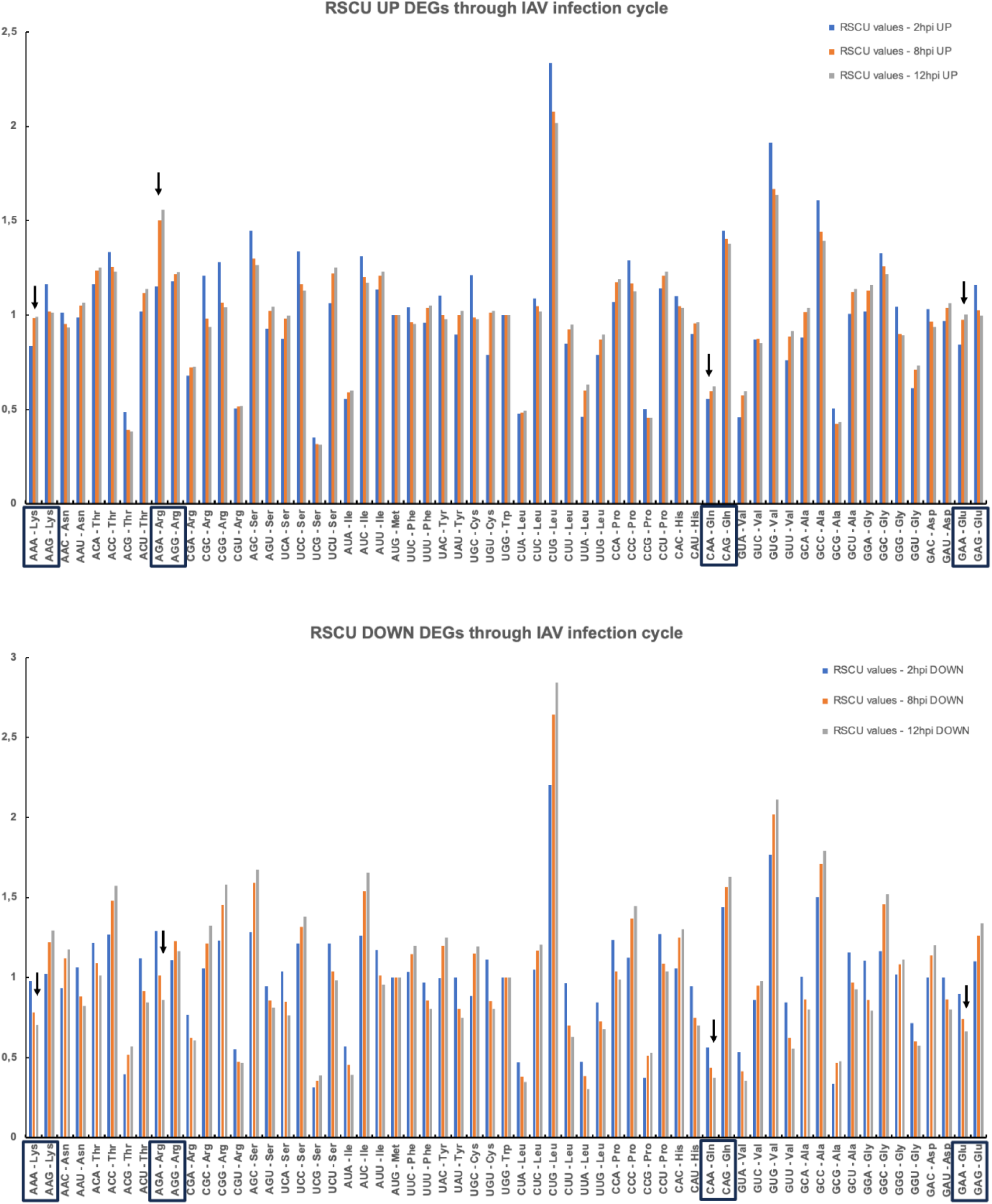
RSCU analysis of up- and down-regulated genes at 2, 8 and 12hpi. Arrows highlight the Lys, Arg, Gln and Glu codons in mixed boxes. Upregulated mRNAs identified at late infection time points (8 and 12 hpi) are preferentially enriched in A-endng codons that are preferentially recognized by tRNAs bearing wobble uridines and mcm5 and mcm5s2 modifications. The opposite trend is observed for downregulat ed mRNAs ate the same IAV infection timepoints.

**Supplementary Figure 4.**
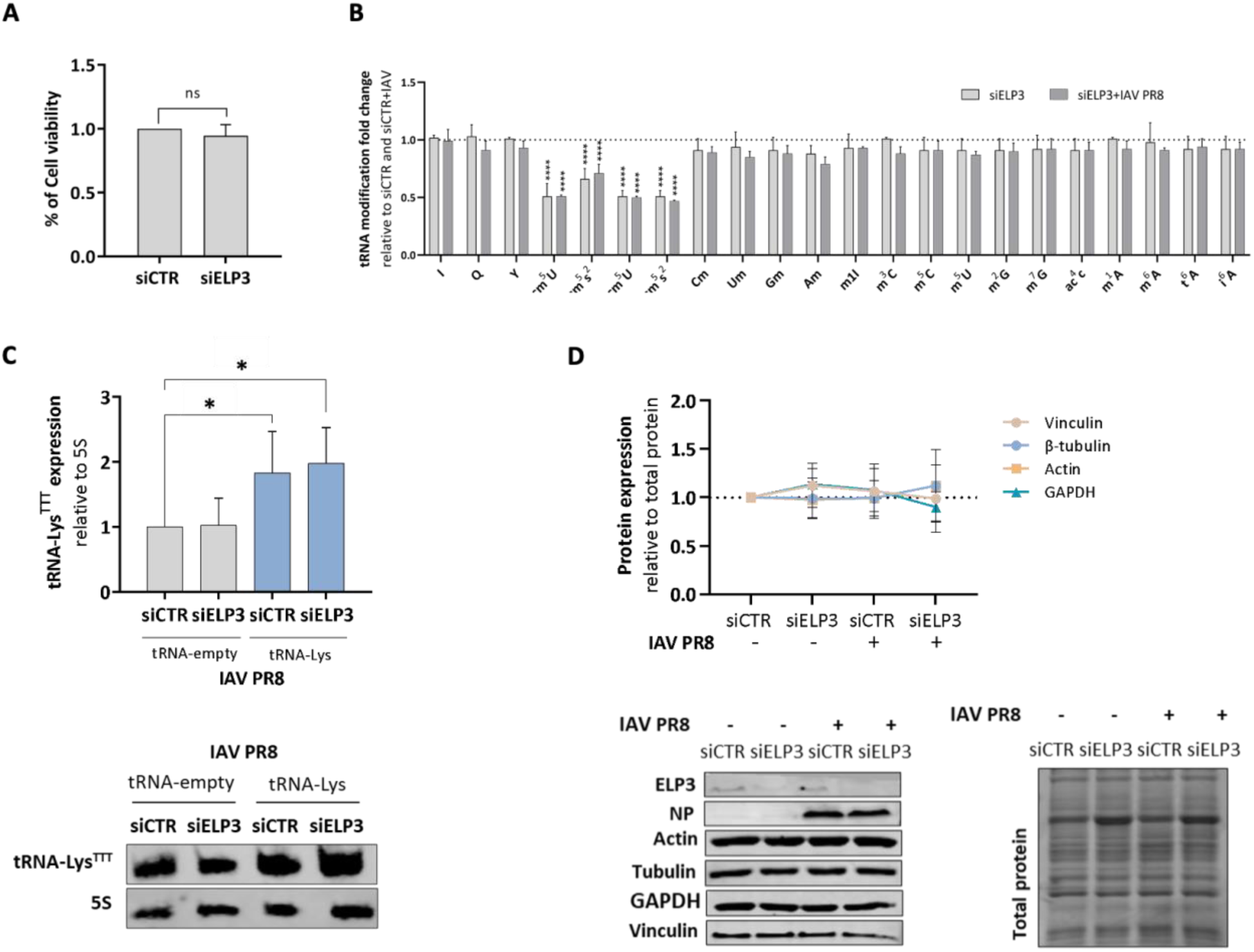
Effect of ELP3 silencing in IAV infected A549 cell lines. (A) Effect of the ELP3 knockdown in A549 cellular viability by the MTT assay. (B) Effect of the ELP3 knockdown in tRNA modification levels by LC-MS/MS. (C)Transfection efficiency of the plasmid encoding the tRNA^Lys^_UUU_ in A549 cells. siCTR- and siELP3- treated cells were transfected with 500 ng of an empty-tRNA vector and the tRNA^Lys^_UUU_ plasmid for 24 hours and then infected with IAV for 8 hours. The levels of tRNA^Lys^_UUU_ were then evaluated by Northen blotting. (D) The levels of several housekeeping proteins were evaluated by western blotting and normalized for total protein levels. A549 cells were infected at a MOI of 3 and all samples were collected at 8 hpi. Data represents the mean of three independent experiments. Bars reflect the mean and error bars reflect standard deviation. All data analysis was done using Student’s unpaired t-test with Welch correction.

**Supplementary figure 5.**
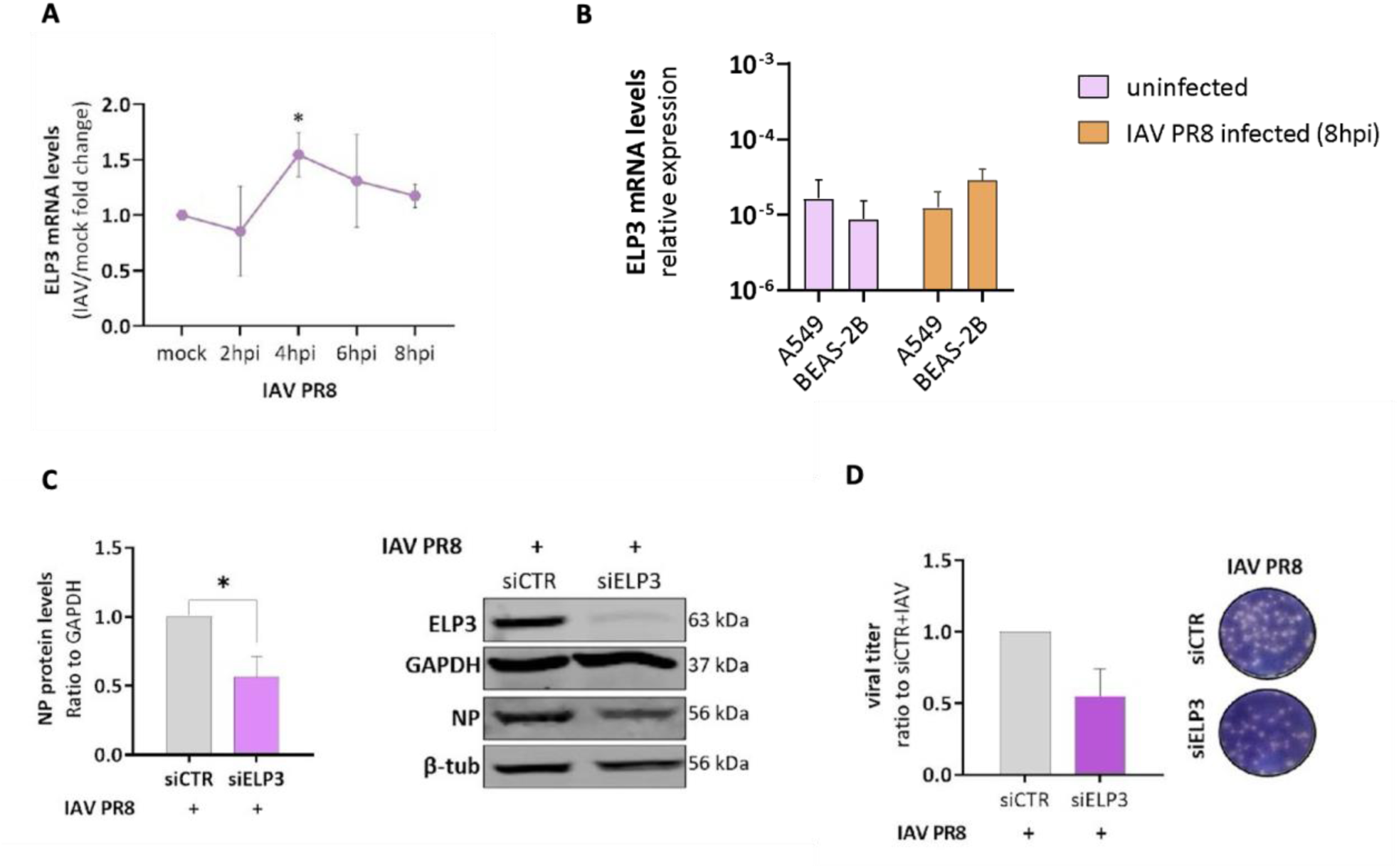
Effect of the ELP3 knockdown in the propagation of IAV in BEAS-2B cell lines. (A) Dynamics of ELP3 abundance upon IAV infection of BEAS-2B cells at different time-points by qRT-PCR. (B) Comparison between basal ELP3 expression levels in BEAS-2B cells and A549 cells. (C) Effect of ELP3 silencing in NP protein levels. (D) Effect of the ELP3 knockdown in IAV PR8 titers by plaque assay. For plaque assays, samples were analyzed at 16hpi. For the remaining experiments, cells were collected at 8hpi. Data represents the mean of at least three independent experiments. Bars reflect the mean and error bars reflect standard deviation. All data analysis was done using Student’s unpaired t-test with Welch correction, *p<0.05 and ****p<0.0001.

**Supplementary Figure 6.**
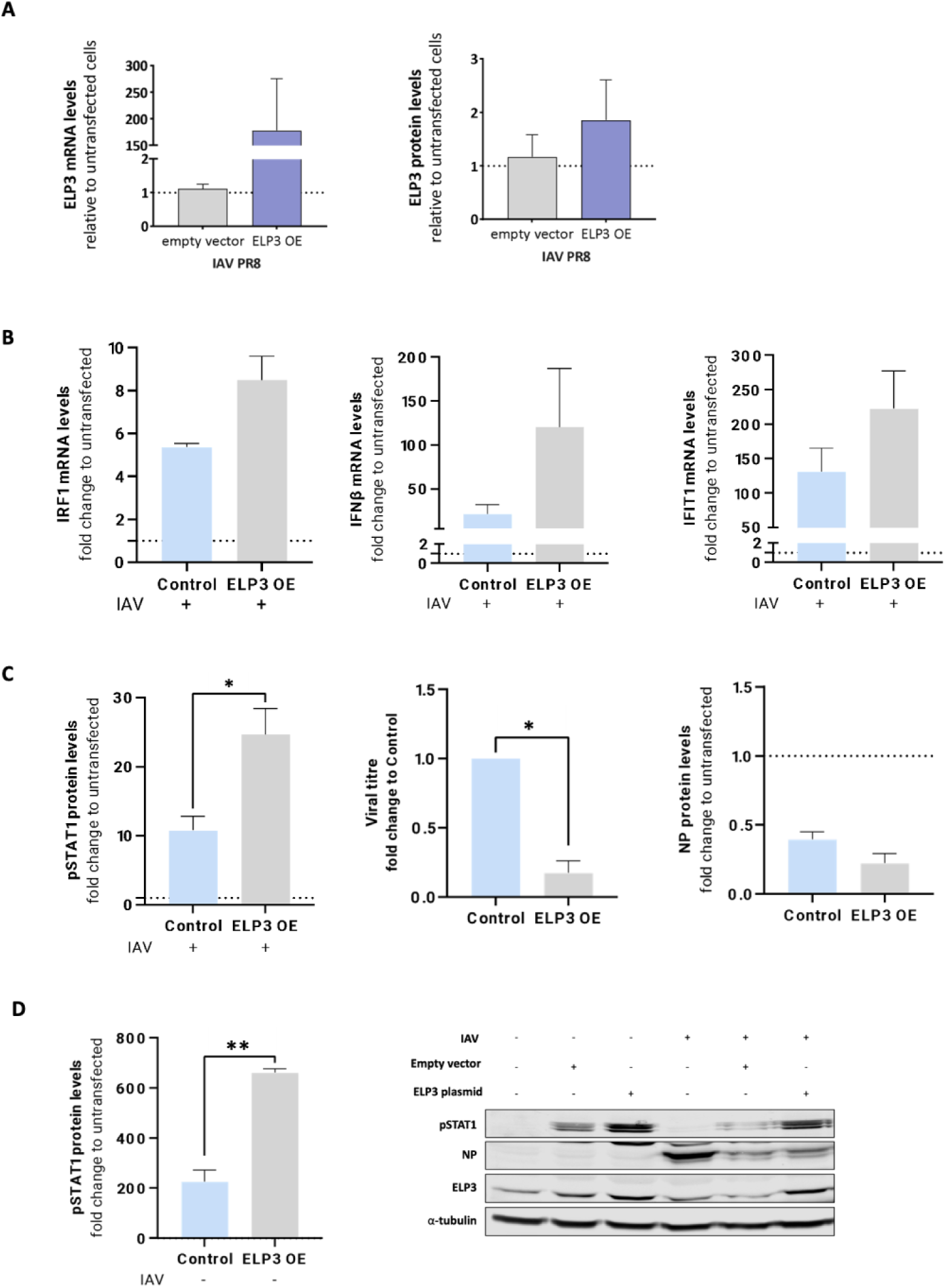
Impact of ELP3 overexpression on innate immune response activation and IAV replication in A549 cells. (A) ELP3 expression levels following ELP3 overexpression in IAV-infected A549 cells. mRNA levels were quantified by qPCR and protein levels by western blotting, and compared with non-transfected cells and cells transfected with an empty vector. Analyses were performed at 8 hours post-infection (hpi). (B) mRNA levels of IRF1, IFNβ, and IFIT1 at 8 hpi in A549 cells overexpressing ELP3 or transfected with an empty vector, relative to non-transfected controls. (C) Quantification of pSTAT1 and viral NP protein levels by western blotting, normalized to α-tubulin and expressed as fold change relative to non-transfected cells (dotted line). Viral particle production was assessed by plaque assay at 16 hpi in supernatants from IAV-infected A549 cells transfected with empty vector or ELP3 plasmid. (D) Basal pSTAT1 levels in non-infected A549 cells overexpressing ELP3, compared with non-transfected and empty vector controls. A549 cells were infected with IAV at MOI = 3. Data are representative of three independent experiments. Statistical significance was determined using an unpaired Student’s *t*-test with Welch’s correction (*p* < 0.05, **p** < 0.01, ***p*** < 0.001). Error bars represent mean ± SD.

**Supplementary figure 7.**
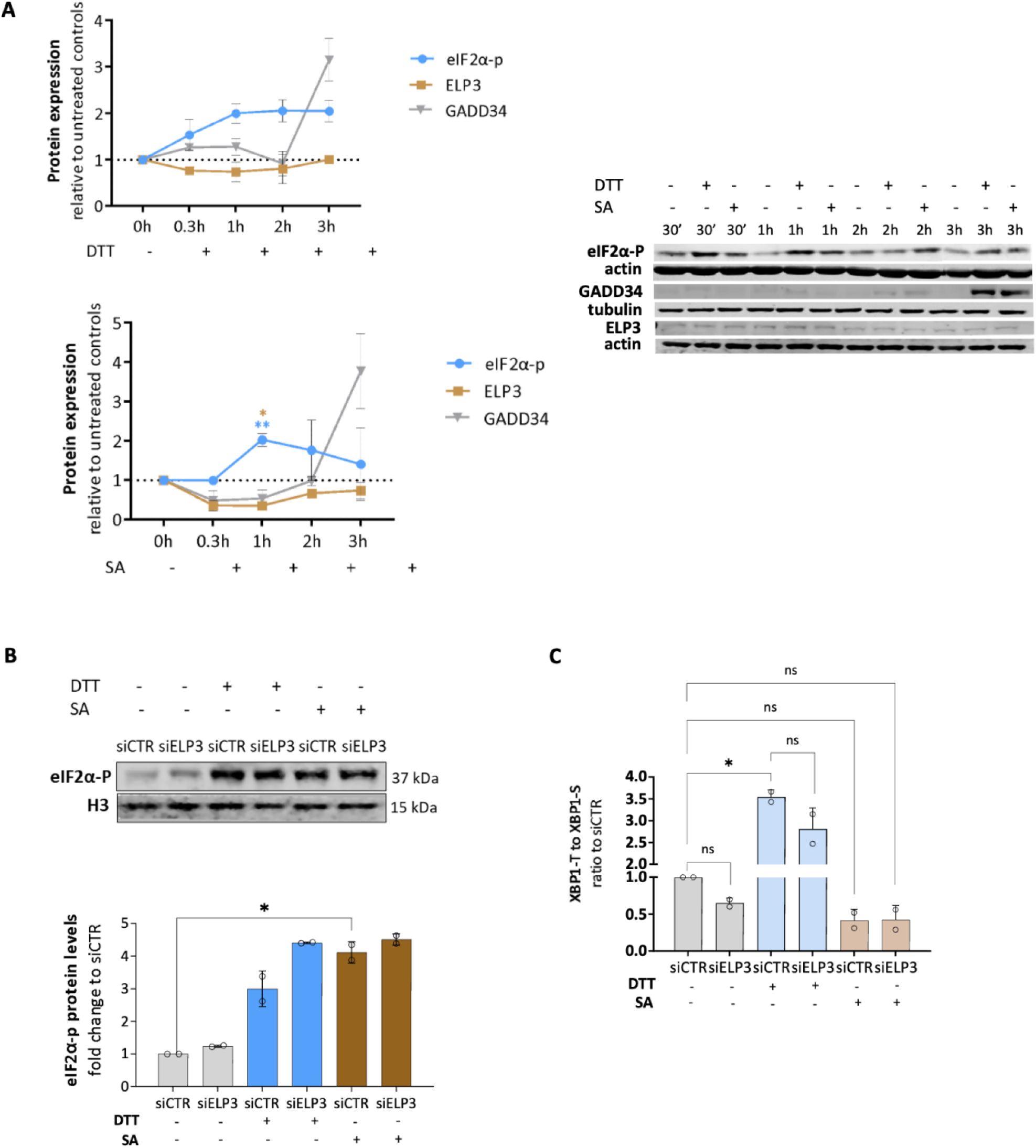
Effect of DTT and SA exposure in ELP3 expression and ER stress in A549 cells. (A) Relative protein expression of eIF2a-P, ELP3 and GADD34 in A549 cells exposed to DTT or SA at different timepoints. Exposure to both stressors elicits eIF2a-P without significant disturbance of ELP3 expression levels. (B) Upon ELP3 silencing there is a slight increase in eIF2a-P levels in cells exposed to both DTT or SA. (C) DTT exposure decreases XBP1 splicing, whereas exposure to SA as the opposite effect. Data represents the mean of at least two independent experiments. Bars reflect the mean and error bars reflect standard deviation. All data analysis was done using Student’s unpaired t-test with Welch correction, *p<0.05 and **p<0.001.

